# The orphan nuclear receptor estrogen-related receptor beta (ERRβ) in triple-negative breast cancer

**DOI:** 10.1101/734632

**Authors:** Aileen I. Fernandez, Xue Geng, Krysta Chaldekas, Brent Harris, Anju Duttargi, V. Layne Berry, Deborah L. Berry, Akanksha Mahajan, Luciane R. Cavalli, Balázs Gyorffy, Ming Tan, Rebecca B. Riggins

## Abstract

**Purpose:** Triple negative breast cancer (TNBC)/ basal-like breast cancer (BLBC) is a highly aggressive form of breast cancer prevalent in African-American (AA) women. We previously reported that a small molecule agonist ligand for the orphan nuclear receptor estrogen-related receptor beta (ERRβ or *ESRRB*) has growth inhibitory and anti-mitotic activity in TNBC cell lines. In this study, we evaluate the association of *ESRRB* mRNA, copy number levels, and protein expression with demographic, clinicopathological, and gene expression features in breast tumor clinical specimens.

**Methods:** *ESRRB* mRNA level expression and clinical associations were analyzed using RNAseq data. Array-based comparative genomic hybridization determined *ESRRB* copy number in AA and Caucasian women. Transcription factor activity was measured using promoter-reporter luciferase assays in TNBC cell lines. Semi-automatic quantification of immunohistochemistry measured ERRβ protein expression on a 150-patient tissue microarray series.

**Results:** *ESRRB* mRNA expression is significantly lower in TNBC/BLBC vs. other breast cancer subtypes. There is no evidence of *ESRRB* copy number loss. *ESRRB* mRNA expression is correlated with the expression of genes associated with neuroactive ligand-receptor interaction, metabolic pathways, and deafness. These genes contain G/C-rich transcription factor binding motifs. The *ESRRB* message is alternatively spliced into three isoforms, which we show have different transcription factor activity in basal-like vs. other TNBC cell lines. We further show that the ERRβ2 and ERRβsf isoforms are broadly expressed in breast tumors at the protein level.

**Conclusions:** Decreased *ESRRB* mRNA expression, and distinct patterns of ERRβ isoform subcellular localization and transcription factor activity are key features in TNBC/BLBC.

## INTRODUCTION

Breast cancer is the most commonly diagnosed cancer in women and is the number two cause of cancer-related death [1]. Breast cancer subtypes are predominantly classified by two methods: immunohistochemistry (IHC) or gene expression. IHC tests for three proteins: estrogen receptor (ER), progesterone receptor (PR), and human epidermal growth factor two (HER2), and based on the expression of these three receptors, patients are classified as having ER+, HER2 overexpressing, or triple negative breast cancer (TNBC). Patients who are diagnosed with TNBC are clinically defined as lacking ER, PR, and HER2 [2]. Gene expression profiling uses a 50-gene panel to determine if a breast cancer is one of 5 intrinsic, or Pam50 subtypes: luminal A, luminal B, HER2-enriched, normal-like and basal-like (BLBC) [3, 4]. In clinic, TNBC and BLBC patients largely converge [5]. TNBC/BLBC is a biologically aggressive subtype of breast cancer with characteristic high genomic instability [6]. It is diagnosed more frequently in African-American (AA) women with much worse prognosis than Caucasian/White (CW) women [7].

Since TNBC patients lack ER and HER2, they are unresponsive to ER and HER2–targeted therapies. Patients must instead be given systemic chemotherapy, which is accompanied by toxic side effects [8]. Due to the aggressive nature of TNBC, it is important to identify new prognostic marker genes capable of defining targets and conversely subtypes within TNBC. Historically, nuclear receptors (NR) have been great targets for cancer treatment, such as the ER in ER+ breast cancer. The NR superfamily consists of many members including orphan nuclear receptors (ONRs) [9] which are defined as lacking any known endogenous ligands. One ONR subgroup contains what are known as the estrogen related receptors (ERR), named for their resemblance to the ER, although they do not bind or respond to estrogen [9]. ERR beta (*ESRRB*, ERRβ), which is alternatively spliced, is one of the first discovered ONRs and is known to have important functions in development [10]. Our lab and others, have previously shown the function of ERRβ in cancer. Our 2016 publication showed that BLBC patients with high *ESRRB* mRNA expression have significantly improved distant-metastases free, and recurrence-free survival in comparison to patients with low expression [11]. Previous publications have shown that BLBC patients overall have significantly lower *ESRRB* mRNA expression in comparison to other breast cancer subtypes [12, 13].

The goal of the present study is to: (1) define *ESRRB* expression levels in breast cancer, specifically in BLBC and TNBC; (2) determine how DNA copy number and protein levels are modulated in various data sets; (3) characterize clinical correlations found in breast cancer. We show that *ESRRB* mRNA expression is significantly lower in BLBC/ TNBC patients and that there is no observed decrease in DNA copy number. ERRβ splice variants have differential transcription factor activity in TNBC cell lines. Lastly, we also show that ERRβ splice variant protein levels are different in breast cancer patient samples depending on the IHC subtype. HER2 and TNBC patients have similar ERRβ protein expression and cellular localization vs. ER+ patients. Our study supports further investigation into the establishment of ERRβ as a TNBC/BLBC therapeutic target or prognostic marker, and as a tool to provide more insights into this aggressive cancer and its mechanism of progression.

### Materials and Methods

#### *In silico* analyses

RNA-sequencing (RNAseq) data was obtained from two publically-available datasets sets: the Sweden Cancerome Analysis Network – Breast (SCAN-B) [14, 15] and The Cancer Genome Atlas (TCGA) [16]. Gene expression profiles of GSE96058 (https://www.ncbi.nlm.nih.gov/geo/query/acc.cgi?acc=GSE96058) were downloaded from Gene Expression Omnibus (GEO) database for the SCAN-B data analyses. This SCAN-B cohort contained 3273 samples (136 were replicates) analyzed by Illumina paired-end RNAseq and expression estimation. This data comes from Sweden and country law states that the submitter cannot provide raw sequence data in a public repository as sequencing may contain personally-identifiable information and hereditary mutations. Data was processed as previously described [14, 15] and gene expression data was generated as FPKM (expression measurement +0.1 FPKM followed by log2 transformation). Clinical information for the publically available SCAN-B data was kindly provided by Dr. Lao Saal (Lund University, Sweden). TCGA raw, processed and clinical data were obtained from the GDC legacy archive (https://portal.gdc.cancer.gov) and accessed using *TCGABiolinks*.

Gene expression analysis was performed using Rstudio (version 3.6.0) and *Bioconductor*. Gene expression levels, measured as FPKM (fragments per kilobase of transcript per million), were determined for all breast cancer patients as two datasets: Pam50 subtypes predetermined by the authors, or IHC subtypes parsed out using clinical information. Overall survival was determined using *survminer*. Differentially expressed genes (DEGs) were detected using *edgeR*. Genes with p<0.05 and fold change ≥2 were considered significant (Table S1). Pathway and represented disease analyses were performed on overexpressed DEGs using KEGG pathway analysis [17, 18].

For SCAN-B data, cox regression was used to determine correlations between *ESRRB* expression and clinical characteristics. For promoter analysis, *Bioconductor* was used to isolate DEGs found within annotated genome UCSC, hg19 and search the promoter region of these genes, −4000 to +500 bp from the transcription start site for enriched short, ungapped, redundant sequences of up to 8 base pairs using Discriminative Regular Expression Motif Elicitation (DREME, [19]). The primary set of sequences was shuffled to create a control set. Significantly overrepresented motifs were determined using Fisher’s Exact test. We then used TomTom [20] to compare our overrepresented motifs to the publically-available JASPAR CORE database [21]which provides the binding preferences of a large database of known transcription factors. Source code for SCAN-B and TCGA analyses can be found at https://github.com/RigginsLabGU/Rmarkdown/blob/master/SCANB%20analysis.Rmd and https://github.com/RigginsLabGU/Rmarkdown/blob/master/TCGA%20analysis.Rmd

#### Array comparative genomic hybridization

The 106-patient cohort of Caucasian (CW) and African American (AA) patients with TNBC and non TNBC (NTN) was collected and processed as previously described by Sugita, *et al* [22]. Copy number was determined from *ESRRB* probes on the Agilent SurePrint aCGH platform. Log_2_ intensity >3 was defined as amplification, ≥/ ≤ 0.25 were defined as copy number gain and loss respectively, and ≥3 was defined as deletion. PRISM 8.0 (Graphpad, San Diego, CA) was used for all statistical analyses of copy number and clinical demographic information.

#### Cell Culture

HCC1806 breast cancer cells were purchased from ATCC (Manassas, VA). MDA-MB-453 breast cancer cells were a gift from Dr. Anna Riegel (Lombardi Comprehensive Cancer Center (LCCC). BT549, MCF7, and MCF10A breast cancer cells were obtained from the LCCC Tissue Culture Shared Resource. Cells were routinely tested for *Mycoplasma spp*. and tested negative and fingerprinted using 9 standard STR loci and Y chromosome-specific amelogenin to verify authenticity. All cells were maintained in a humid carbon dioxide (CO_2_) incubator, 95% air: 5% CO_2_. HCC1806 and MDA-MB-453 cells were grown in improved minimal essential media (IMEM; Life Technologies, Grand Island, NY) supplemented with 10% heat-inactivated fetal bovine serum (FBS, purchased from LCCC Tissue Culture Shared Resource). BT549 cells were cultured in IMEM with 10% FBS and 10 μg/mL insulin (purchased from Life Technologies, Grand Island, NY).

#### Plasmids and transfection

The psG5 empty vector, ERRβsf (murine ERRβ, >90% homology to human ERRβ, Addgene #52188), ERRβ2 (Addgene #52186), and ERRβ-Δ10 (Addgene #52187) constructs have all been published previously [11, 23]. The ERRE-luciferase (Addgene #37851) and p21-luciferase (Addgene #21723) have been previously described [23]. Plasmids were introduced using Mirus TransIT-X2 Dynamic Delivery System (Mirus Bio LLC; MIR600Q) according to manufacturer’s instructions.

#### Dual-luciferase promoter-reporter assays

Cells were seeded into 24-well plastic tissue culture dishes at 150,000 cells per well on day 0. On day 1, cells were transfected using 500 ng DNA/ well (137 ng receptor; 360 ng luciferase reporter plasmid; 3 ng Renilla control) for 24 hours. On day 2, media containing the transfection complexes was removed and fresh media was added to the cells. On day 3, at 48 hours, cells were harvested for luciferase assay (https://www.promega.com/products/reporter-assays-and-transfection/reporter-assays/dual_luciferase-reporter-assay-system/) according to the manufacturer’s instructions. Luciferase activity was normalized to Renilla. All experiments were performed 3-5 times.

#### Western blotting and antibodies

Lysate collected for dual-luciferase activity assay were run on 4-12% poly acrylamide gels using electrophoresis for 90 minutes. After protein was transferred to nitrocellulose membranes, the membranes were blocked for one hour in 5% nonfat dry milk in Tris-Buffered Saline with Tween-20 (TBST) and probed overnight at 4°C with ERRβ #PP-H6707-00 (cl.07) 1:500. All membranes were then re-probed with loading control β-tubulin at 1:10:000 for 1 hour at room temperature or overnight at 4°C.

Horseradish peroxidase enzyme-conjugated anti-mouse whole immunoglobulin (IgG) secondary antibody (GE #NXA931 Buckinghamshire, U.K.) was used the following day at 1:5000 for 1 hour at room temperature. The membranes were then imaged for enhanced chemiluminescence (ECL, Denville Scientific, Holliston, MA) on the Amersham Imager 600 (GE Life Sciences).

#### Image analysis and statistical analysis of *in vitro* work

Images and figures were composed using Adobe Photoshop, Illustrator and InDesign. FIJI was used to perform densitometry on imaged blots. Statistical analyses of dual-luciferase activity assay and western blot densitometry were done using PRISM 8.0 (Graphpad, San Diego, CA).

#### Invasive Ductal Carcinoma Breast Cancer TMA series

The Histopathology and Tissue Shared Resource (HTSR) at Georgetown University Medical Center’s Lombardi Comprehensive Cancer Center (LCCC) constructed the Invasive Ductal Carcinoma Tissue Microarrays (TMA) series. The cohort consists of 150 breast cancer patients distributed as 50 patients each on a series of 3 TMAs. All patients are research-consented through the HTSR, the Survey, Recruitment and Biospecimen Shared Resource (SRBSR), and/or Indivumed groups under the following respective Georgetown University Medical Center IRB protocols: 1992-048, Pr0000007, and 2007-345. The 50 patients for each TMA were grouped based on the following molecular subtypes: ER alpha positive TMA (>10% ER alpha positivity, PR+/-, HER2 negative), HER2 positive TMA (ER+/-, PR+/-) and Triple Negative Breast Cancer (TNBC) TMA. The TMAs were stained for ER alpha, PR, HER2, Ki67 and panCytokeratin in a single multiplexed assay. High resolution images are available for all stained cores as well as quantification of percentage positive for ER, PR, and Ki67 and threshold analysis for HER2 positivity. 2 cores for each patient are available side by side on the TMA. Each TMA has matching biological and immunological controls including tonsil, spleen, testis, reduction mammoplasty (benign breast), placenta and 6 well characterized breast cancer cell lines. Inclusion criteria for the TMA were female patients diagnosed with invasive ductal carcinoma with at least 3 years clinical follow up (majority with 5-year follow up, or otherwise deceased) with a primary breast cancer surgical resection at MedStar Georgetown University Hospital (MGUH) between 2004-2014. Exclusion criteria were male patients, patients diagnosed with ductal carcinoma in situ only or lobular carcinoma, known BRCA or familial mutation carriers, or evidence of neoadjuvant therapy. Racial distribution of the patients was 64% White/Caucasian, 26% Black/African American, 10% other or unknown, allowing for support for projects analyzing racial disparities. Clinical, treatment and follow up data were retrieved by the Innovation Center for Biomedical Informatics (ICBI) from the MGUH Cancer Registry. Pathology data was manually extracted from the original surgical pathology reports. All demographic, clinical, pathology, and available follow up data were de-identified and uploaded into a REDCap database for query and analysis. High resolution images of hematoxylin and eosin stained cores are available for all cores on the TMA.

#### Immunohistochemical staining

IHC staining of breast cancer tissue was performed for ERRβ. Five micron sections from formalin fixed paraffin embedded tissues were de-paraffinized with xylenes and rehydrated through a graded alcohol series. Heat induced epitope retrieval (HIER) was performed by immersing the tissue sections in Target Retrieval Solution, Low pH (DAKO) in the PT Link (DAKO). IHC staining was performed using the VectaStain Kit from Vector Labs according to manufacturer’s instructions. Briefly, slides were treated with 3% hydrogen peroxide, avidin/biotin blocking, and 10% normal goat serum and independently exposed to primary antibodies for ERRβsf-cl.07, 1:150, 1:240 (R&D systems, #PP-H6707-00) and ERRβ2-cl.05, 1:150 (R&D systems, #PP-H6705-00) for 1 hour at room temperature. Slides were exposed to appropriate biotin-conjugated secondary antibodies (Vector Labs), Vectastain ABC reagent and DAB chromagen (Dako). Slides were counterstained with Hematoxylin (Fisher, Harris Modified Hematoxylin), blued in 1% ammonium hydroxide, dehydrated, and mounted with Acrymount. Control tissues with the primary antibody omitted were used as negative controls.

#### Scanning and Analysis using Vectra3

Stained slides were scanned using the Vectra3 Multi-Spectral Imaging Microscope with Vectra and Phenochart software (Perkin Elmer). Every available TMA core was imaged as a 3×3 image to capture almost the entire core in one image. The scanned images were analyzed in inForm software version 2.3. Nuclear versus cytoplasmic staining was differentiated. The individual phenotypes were combined in Microsoft Excel to identify cell phenotypes as ERRβ positive. An average of 11,000 cells were counted per TMA core.

#### Statistical analysis of tissue microarray data

To prepare for statistical analysis, individual cores were manually examined and matched to their corresponding coordinates defined by Vecta3 software. Any cores missing more that 50% of tissue due to adipose tissue or folding were omitted from final analysis. The two cores from each patient were individually scored then averaged to determining the 50 scores per TMA slide used in the analyses.

ERRβ-clone 07 (recognizes ERRβ2), ERRβ-clone 05 (recognizes ERRβsf), and their ratio, as well as demographic variables were summarized by using mean (standard deviation, sd) and median (interquartile range, IQR) for continuous variables and frequency and percentage for categorical variables. Kruskal-Wallis tests were used to test whether median ERRβ2 and ERRβsf expression, and their ratio, were significantly different among the three IHC receptor subtypes. In order to assess whether the expression of each isoform was significantly different among the three IHC subtypes while considering lymph node status, race, and age, and if there was any interaction between IHC subtype and demographic variables, we first made a logit transformation of ERRβ-clone 07 and ERRβ-clone 05 which achieved approximate normality. Then two-way ANOVA was used for analysis following ANOVA procedure. If a significant difference was observed, pairwise comparisons were performed by Dwass, Steel, Critchlow-Fligner multiple comparison procedure (DSCF).

Nuclear staining and cytoplasmic staining were summarized by IHC subtype using mean (sd) and median (IQR). Spearman’s correlation coefficient was calculated to measure the association between the nuclear and cytoplasmic staining in each IHC subtype.

All tests were two-sided at a significant level 0.05. No method had been used for adjusting multiple comparisons. All analyses were performed using SAS 9.4 and Rstudio (Version 0.99.902) software.

## RESULTS

### Low *ESRRB* mRNA expression is associated with shorter overall survival

We previously published that high expression of *ESRRB* mRNA is associated with improved recurrence-free and distant-metastasis free survival in a merged cohort of BLBC patients from multiple independent studies [11, 24]. Here, we analyzed the association of *ESRRB* mRNA expression with overall survival by analyzing Illumina HT 12 gene array data from the Molecular Taxonomy of Breast Cancer International Consortium (METABRIC) study [25]. Importantly, patients selected for this analysis were systemically untreated, providing a more compelling link between *ESRRB* expression and clinical outcome that is not confounded by treatment effect. We found that low *ESRRB* expression (defined as below median) is associated with significantly shorter overall survival (OS) only in BLBC patients (Fig. 1).

**Fig. 1.**
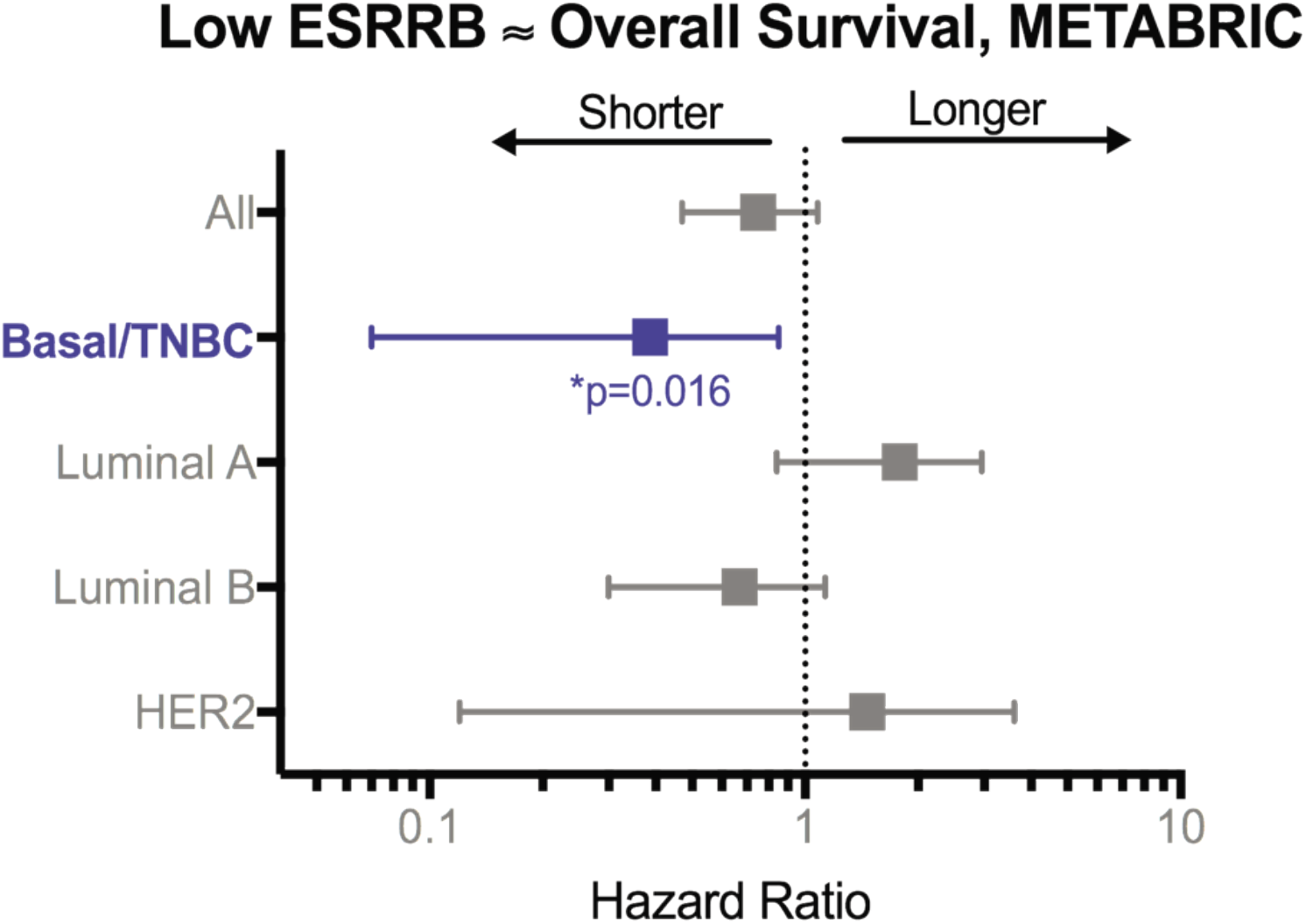
ESRRB association with overall survival (OS) in women with systemically untreated breast cancer. Hazard ratios, 95% confidence intervals, and log-rank p value for low *ESRRB* expression (METABRIC RNAseq data, below median) relative to OS.

### *ESRRB* mRNA expression levels are significantly decreased in BLBC and TNBC, but not associated with age, grade, or lymph node status

We used two large, publicly-available RNAseq data sets –SCAN-B [14, 15] and The Cancer Genome Atlas (TCGA) [16] – to analyze *ESRRB* mRNA expression in association with demographic, clinicopathologic, and gene expression features. *ESRRB* expression as FPKM was measured in both data sets in patients stratified by either IHC or Pam50 breast cancer subtypes for analysis (Fig. 2). In SCAN-B, *ESRRB* expression was significantly lower in BLBC patients compared to luminal A and normal-like patients (Fig. 2a). *ESRRB* expression was also significantly lower in TNBC patients vs. ER+ and ER+/HER2+ patients (Fig. 2b). In TCGA data, we confirmed the previously published findings of Garattini, *et al*. [13] and found that BLBC patients had significantly lower *ESRRB* expression compared to luminal A patients (Fig. 2c). Like the SCANB cohort, TCGA-TNBC patients also had significantly lower *ESRRB* expression compared to ER+ and ER+/HER2+ patients (Fig. 2d).

**Fig. 2.**
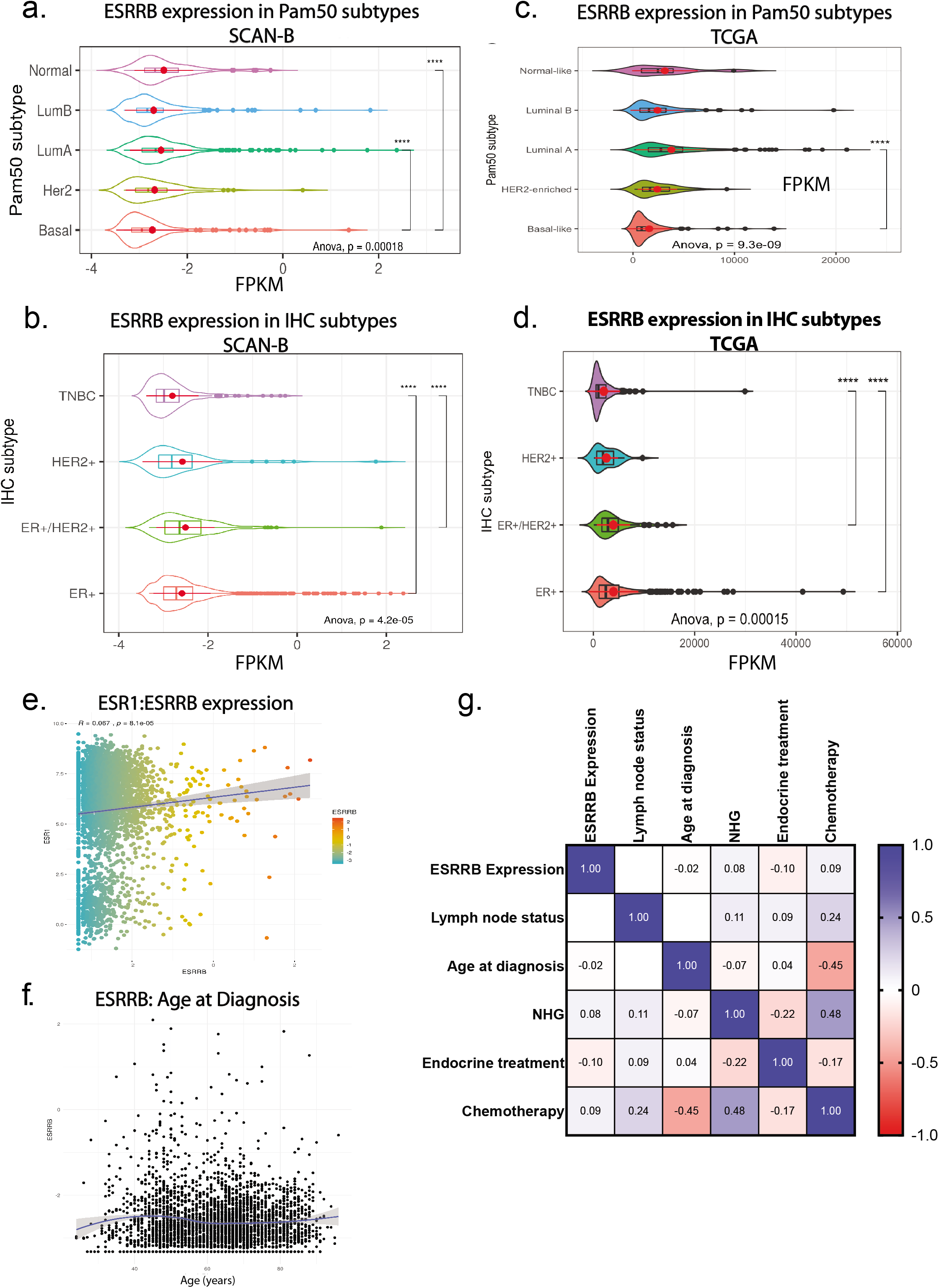
ESRRB mRNA expression in SCAN-B and TCGA data sets. Analysis of RNAseq data from SCAN-B **(a, b, e, f, g)** and TCGA **(c, d)** data. Kruskal-Wallis one-way ANOVA and Tukey multiple comparisons of means with 95% family-wise confidence level. **a-d**. *ESRRB* mRNA FPKM levels by PAM50 subtype in SCAN-B **(a)** and TCGA **(c)** data. *ESRRB* mRNA levels by IHC subtype in SCAN-B **(b)** and TCGA (d) data. Shown as mean ± standard deviation, p < 0.0001 **** **e-f.** Correlations in SCAN-B dataset. **e.** Correlation of *ESR1* to *ESRRB* mRNA expression **f.** *ESRRB* mRNA expression by age **g.** Correlation of *ESRRB* high and low expression, frequency of node status, age at diagnosis, NHG, endocrine treatment, and chemotherapy in all breast cancer patients in dataset. Spearman correlation. White spaces were not significantly different. Light to dark colors, correlation observed and statistically significant, p <0.001

SCAN-B is a 3273-patient data set collected in Sweden starting in 2010 [14, 15]. Biospecimens were collected and RNAseq was performed on biopsy cores as previously described [14]. Comprehensive clinical and demographic information was also collected, allowing for analysis of *ESRRB* association with these data for both the Pam50 and IHC subtypes. Analysis of Pam50 subtype revealed BLBC patients were significantly younger than luminal A, luminal B, and HER2 patients (Fig. S1a), a characteristic commonly found in BLBC patients, while analysis by IHC subtype showed that HER2 patients were significantly younger that ER+ patients (Fig. S1b).

We also assessed the correlation between *ESRRB* high and low mRNA expression and clinical characteristics. Across the entire SCAN-B cohort, there was a weak but statistically significant positive correlation between estrogen receptor (*ESR1)* and *ESRRB* mRNA expression (p = 8.1 e-05, Fig. 2e). There was no significant correlation between ESRRB expression and age (Fig. 2f). Filtered data sets were split into tertiles with the top third considered *ESRRB* “high” expression and the lower third considered “low” expression. Analysis within the Pam50 and IHC subtypes showed that there was no significant difference in age between *ESRRB* high and low patients within the Pam50 subtypes (not shown). There also were no significant differences observed in lymph node status, Nottingham grade (NHG), endocrine treatment or chemotherapy treatment between *ESRRB* high and low patients. However, receipt of chemotherapy was negatively correlated with age at diagnosis (p < 5e^-107^) and positively associated with Nottingham grade (NHG, p < 6.7e^-127^), which is broadly reflective of clinical management decisions made in the treatment of younger women and higher grade disease (Fig. 2g) [26].

OS was assessed comparing *ESRRB* high and low patients (upper vs. lower tertiles) in all breast cancer, BLBC, and TNBC patients from TCGA and SCAN-B datasets. There was no statistically significant difference in OS found for *ESRRB* high vs. low patients in SCAN-B or TCGA data (Fig. S1c-S1h). This is likely because in SCAN-B, >80% of the patient population remain alive, while in the TCGA data set filtering out TNBC and BLBC patients leaves a low number of patients with an event (n = 123/1098) decreasing the power of these analyses. Though not statistically significant, BLBC patients in the TCGA data set with high *ESRRB* expression did trend towards better OS than *ESRRB* low patients (hazard ratio = 2.13). Overall, these data confirm lower *ESRRB* expression in TNBC and BLBC patients from two large-scale cohorts but identify no statistically significant associations between *ESRRB* expression and age, grade, lymph node status, or treatment.

### *ESRRB* does not have copy number loss in TNBC

Next, we sought to determine if the observed decrease in *ESRRB* mRNA expression found in TNBC/BLBC may be due to copy number loss. We approached this by analyzing an independent, 106-patient cohort of Caucasian (CW) and African American (AA) patients with TNBC and non TNBC (NTN) using array comparative genomic hybridization (aCGH) as previously described by Sugita, *et al* [22]. *ESRRB* copy number was calculated and defined as deletion, loss, gain, or amplification, and demographic information was analyzed for associations with *ESRRB* copy number changes (Fig. S2a-e). Fisher’s exact test showed that in our cohort, patients with TNBC had significantly higher grade than non-triple negative (NTN) patients at diagnosis (p < 0.0001), indicating that our cohort is representative of the TNBC population (Fig. 3a). There were no significant differences observed between CW and AA patients with TNBC and NTN in relation to the total number of copy number alterations. However, > 75% of all patients showed copy number gain rather than loss at the *ESRRB* locus, 14q24.3 (Fig. 3b).

**Fig. 3.**
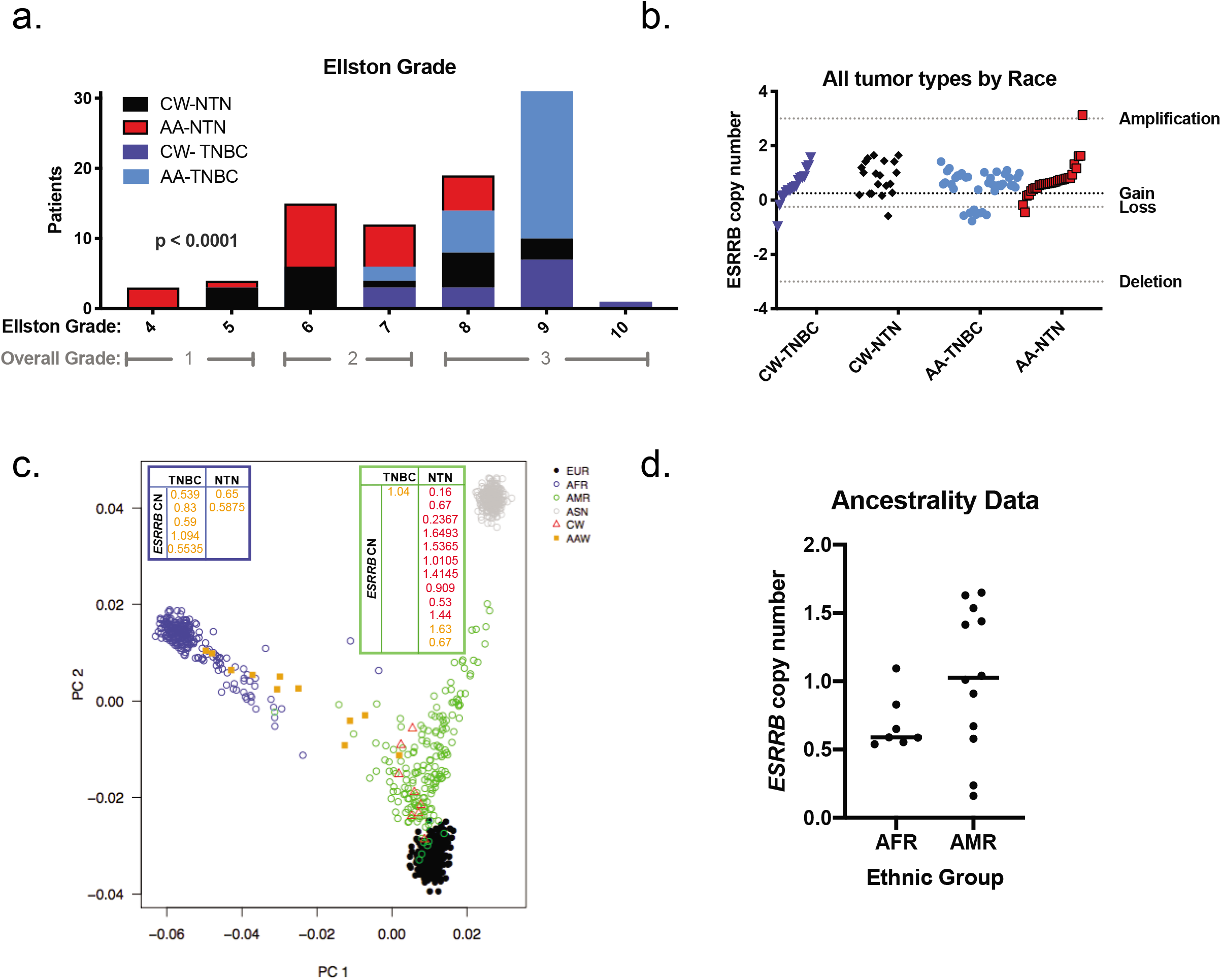
*ESRRB* copy number in breast tumor specimens. **a.** Women with TNBC have higher grade than women with nonTNBC, Fisher’s exact test p<0.0001 **** **b.** Copy number determined from *ESRRB* probes on the Agilent SurePrint aCGH platform. Log_2_ intensity >3=amplification, >0.25=gain, ≤0.25=loss, ≥3=deletion. Shown by Caucasian (CA) vs. African American (AA) women, TNBC vs nonTNBC patients. Fisher’s exact test, all tumors, CA vs AA, n.s. **c, d.** Ancestrality. SNP chip Illumina Infinium QC Array (Illumina Inc., CA) looking at ~3,000 ancestral informative markers in a portion of the Caucasian American (CW) and African American (AA) patients included in the ESRRB copy number study. Four super populations were identified: **European (EUR)**, African (AFR), Ad mixed American (AMR), and East Asian (ASN). **d.** *ESRRB* copy number in self-identified AA or CW redistributed as 1000s genomes project AFR descent or AMR descent. Mann-Whitney, n.s

Since self-reported race and ethnicity data are not always concordant with genotyping analyses, we also assessed ancestry informative markers (AIMs) and the distribution of *ESRRB* copy number in a small subset of our cohort [25, 27]. This 20-patient subset included 6 TNBC patients and 14 NTN patients. Principal component analysis (PCA) of 3000 AIMs overlaid with self-reported ethnicity of our cases showed separation of distinct ancestral populations [22] when merged with two populations from the 1000 Genomes Project: African descent (AFR) and ad mixed American (AMR) (Fig. 3c). The mean rank for *ESRRB* was markedly lower in the AA, but there was no statistically significant difference in the average *ESRRB* copy number in breast tumors from the AA women (clustered with the AFR group) versus the CW (clustered with the AMR group) (Fig. 3d).

### Differentially expressed genes in *ESRRB* high versus *ESRRB* low patients

We next identified DEGs in *ESRRB* mRNA high versus low patients to determine what other genes are co-modulated with *ESRRB* expression. First, we compared *ESRRB* high and low patients (upper and lower tertiles, respectively) and assessed genes that were high in ESRRB-high patients (left/ pink) and low in ESRRB-high patients (right/ purple) (≥2-fold up-or down-regulated, p < 0.05, Fig. 4a-d; Table S1). We determined the overlap of DEGs between data sets and between BLBC and TNBC, finding the most overlap within datasets (Fig. 4e, Fig. S3a, b). Prior studies report that greater than 80% of TNBC patients are also BLBC [5] and our analysis of the SCAN-B and TCGA data sets shows a similar overlap (Fig. S3c,d). KEGG pathway analysis [17, 18] found enrichment of genes associated with neuroactive ligand-receptor interaction (fold change ≥ 3.65) and metabolic pathways (fold change ≥ 2.34) overexpressed in *ESRRB* high patients. Three fourths of the DEGs lists included deafness as one of the disease associations. Mutations in *ESRRB* are linked to a hereditary autosomal recessive hearing disorder [28, 29] (Figure 4f).

**Fig. 4.**
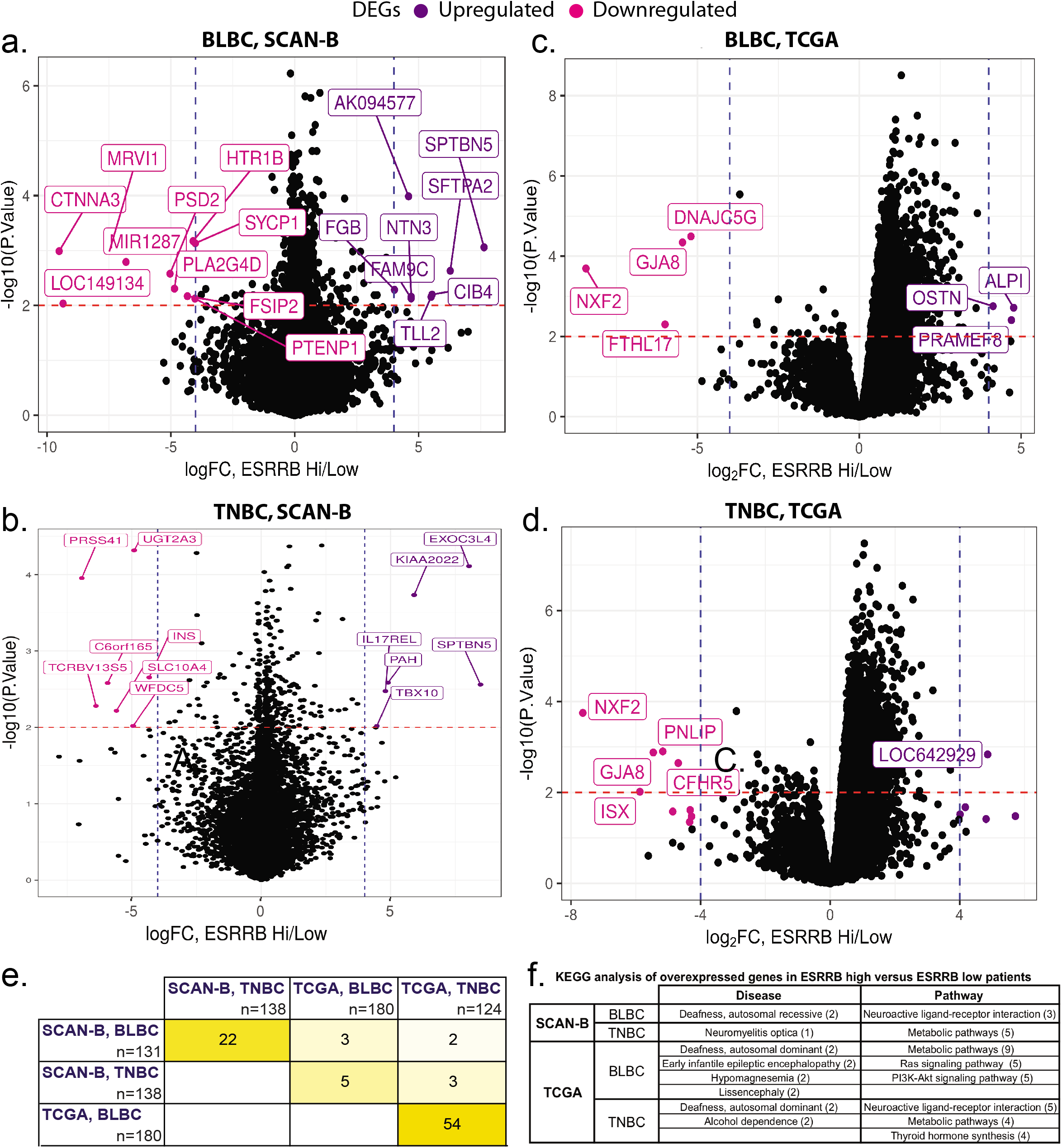
Differentially expressed genes (DEGs) associated with *ESRRB* expression in SCAN- and TCGA data sets. DEG analysis in patients with BLBC **(a, c)** and TNBC **(b, d)**. Shown are DEGs that are high in ESRRB high patients, and low in ESRRB high patients in SCANB data, p<0.01, fold change (FC) > 4. **e.** Heat map depicting DEG overlap between the four analyses shown in a-e. **f.** Top pathways and diseases represented by overexpressed DEGs in *ESRRB* high-versus-low patients.

We then searched for transcription factor binding motifs within the promoter region of the enriched DEGs in the SCAN-B dataset [19]. DEGs in BLBC samples had 39 enriched motifs which aligned to 320 known transcription factors TNBC samples had 40 enriched motifs which aligned to 611 known transcription factors (TOMTOM [20]; JASPAR [21], Fig. S3e). Top matches for both BLBC and TNBC in the SCAN-B dataset were extracted, E<0.05. The top motifs found in BLBC and TNBC were GGCACGTGCC and CCACCGACA, respectively. These motifs aligned to multiple known transcription factors, including basic helix-loop-helix (bHLH) and AP-2 motifs, both characterized as G/C-rich (Table 1).

**Table 1.**
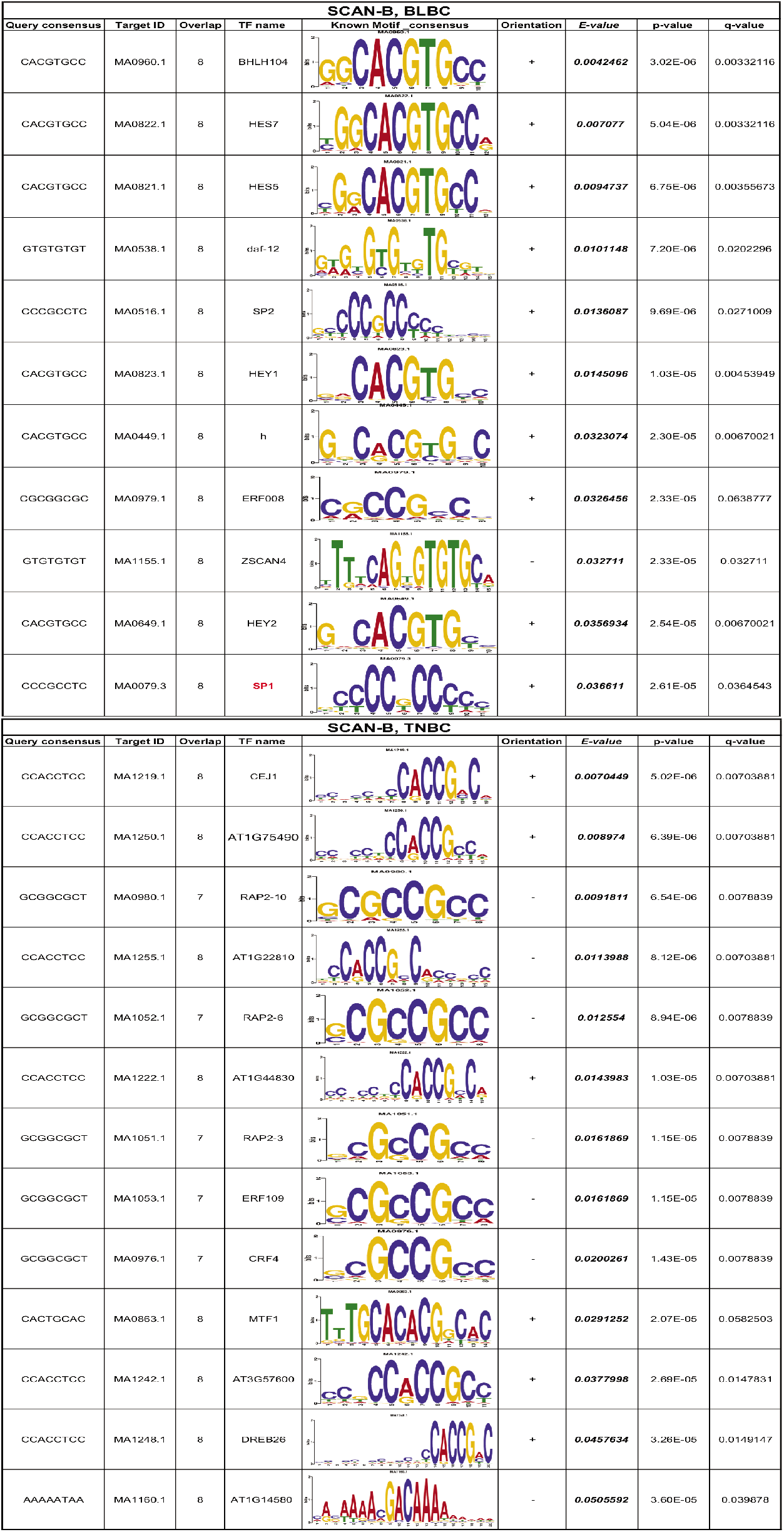
TOMTOM analysis of enriched transcription factor binding motifs. Top motifs from DEGs aligned to JASPAR non-redundant DNA database. Corresponding motifs of known transcription factors are SCAN-B and TCGA data. Determined in BLBC and TNBC

| SCAN-B, BLBC |  |  |  |  |  |  |  |  |
| --- | --- | --- | --- | --- | --- | --- | --- | --- |
| Query consensus | Target ID | Overlap | TF name | Known Motif consensus | Orientation | E-value | p-value | q-value |
| CACGTGCC | MA0960.1 | 8 | BHLH104 |  | + | 0.0042462 | 3.02E-06 | 0.00332116 |
| CACGTGCC | MA0822.1 | 8 | HES7 |  | + | 0.007077 | 5.04E-06 | 0.00332116 |
| CACGTGCC | MA0821.1 | 8 | HES5 |  | + | 0.0094737 | 6.75E-06 | 0.00355673 |
| GTGTGTGT | MA0538.1 | 8 | daf-12 |  | + | 0.0101148 | 7.20E-06 | 0.0202296 |
| CCCGCCTC | MA0516.1 | 8 | SP2 |  | + | 0.0136087 | 9.69E-06 | 0.0271009 |
| CACGTGCC | MA0823.1 | 8 | HEY1 |  | + | 0.0145096 | 1.03E-05 | 0.00453949 |
| CACGTGCC | MA0449.1 | 8 | h |  | + | 0.0323074 | 2.30E-05 | 0.00670021 |
| CGCGGCGC | MA0979.1 | 8 | ERF008 |  | + | 0.0326456 | 2.33E-05 | 0.0638777 |
| GTGTGTGT | MA1155.1 | 8 | ZSCAN4 |  | - | 0.032711 | 2.33E-05 | 0.032711 |
| CACGTGCC | MA0649.1 | 8 | HEY2 |  | + | 0.0356934 | 2.54E-05 | 0.00670021 |
| CCCGCCTC | MA0079.3 | 8 | SP1 |  | + | 0.036611 | 2.61E-05 | 0.0364543 |

| SCAN-B, TNBC |  |  |  |  |  |  |  |  |
| --- | --- | --- | --- | --- | --- | --- | --- | --- |
| Query consensus | Target ID | Overlap | TF name | Known Motif consensus | Orientation | E-value | p-value | q-value |
| CCACCTCC | MA1219.1 | 8 | CEJ1 |  | + | 0.0070449 | 5.02E-06 | 0.00703881 |
| CCACCTCC | MA1250.1 | 8 | AT1G75490 |  | + | 0.008974 | 6.39E-06 | 0.00703881 |
| GCGGCGCT | MA0980.1 | 7 | RAP2-10 |  | - | 0.0091811 | 6.54E-06 | 0.0078839 |
| CCACCTCC | MA1255.1 | 8 | AT1G22810 |  | - | 0.0113988 | 8.12E-06 | 0.00703881 |
| GCGGCGCT | MA1052.1 | 7 | RAP2-6 |  | - | 0.012554 | 8.94E-06 | 0.0078839 |
| CCACCTCC | MA1222.1 | 8 | AT1G44830 |  | + | 0.0143983 | 1.03E-05 | 0.00703881 |
| GCGGCGCT | MA1051.1 | 7 | RAP2-3 |  | - | 0.0161869 | 1.15E-05 | 0.0078839 |
| GCGGCGCT | MA1053.1 | 7 | ERF109 |  | - | 0.0161869 | 1.15E-05 | 0.0078839 |
| GCGGCGCT | MA0976.1 | 7 | CRF4 |  | - | 0.0200261 | 1.43E-05 | 0.0078839 |
| CACTGCAC | MA0863.1 | 8 | MTF1 |  | + | 0.0291252 | 2.07E-05 | 0.0582503 |
| CCACCTCC | MA1242.1 | 8 | AT3G57600 |  | + | 0.0377998 | 2.69E-05 | 0.0147831 |
| CCACCTCC | MA1248.1 | 8 | DREB26 |  | + | 0.0457634 | 3.26E-05 | 0.0149147 |
| AAAAATAA | MA1160.1 | 8 | AT1G14580 |  | - | 0.0505592 | 3.60E-05 | 0.039878 |

### ERRβ isoform transcription factor activity differs across cellular models of TNBC molecular subtypes

A key challenge in defining bona fide *ESRRB* target genes amongst DEGs from high dimensional data sets is that ERRβ, at the protein level, is expressed as at least three distinct isoforms: ERRβsf, ERRβ2, ERRβ-Δ10 [30, 31]. These isoforms are produced via alternative splicing in which a genetic alteration at intron-exon boundaries, or splice sites, are altered at the mRNA level resulting in different proteins [32]. Two of these isoforms – ERRβsf and ERRβ2 – have opposing functions in regulating cell cycle progression [11], while less is known about ERRβ-Δ10 due to a lack of validated antibodies to detect this isoform at the protein level [33]. We first established basal level of two ERRβ isoforms - ERRβ2 and ERRβsf - in a panel of TNBC cell lines reflective of diverse TNBC subtypes. TNBC subtypes were initially defined as a step to further personalize medicine. There are 6 distinct TNBC subtypes based on gene expression and ontology [34, 35]. Cell lines representing 5 of the 6 subtypes, as well as nontransformed mammary epithelial cells MCF10A and estrogen-receptor positive cells MCF7, were probed for ERRβ2 and ERRβsf using two monoclonal antibodies we have previously characterized as selective for each isoform [11, 23]. TNBC cell lines had a trend toward lower ERRβ protein levels in comparison to the MCF10As and MCF7s (Fig. 5a-c, Figure S4a, b). HCC1806s (basal-like 2, BL2) had the lowest ERRβ2:ERRβsf ratio while MDA-MB-453s (luminal androgen receptor, LAR) had the highest ratio, though these differences did not reach statistical significance. Consistent with this, assessment of TNBC subtypes at the mRNA level in SCAN-B tumor also showed no significant differences in *ESRRB* expression between the subtypes (Fig. S4c). Analysis of publicly available gene expression data in cell lines showed similar ERRβ2 mRNA expression (Figure S4d, e).

**Fig. 5.**
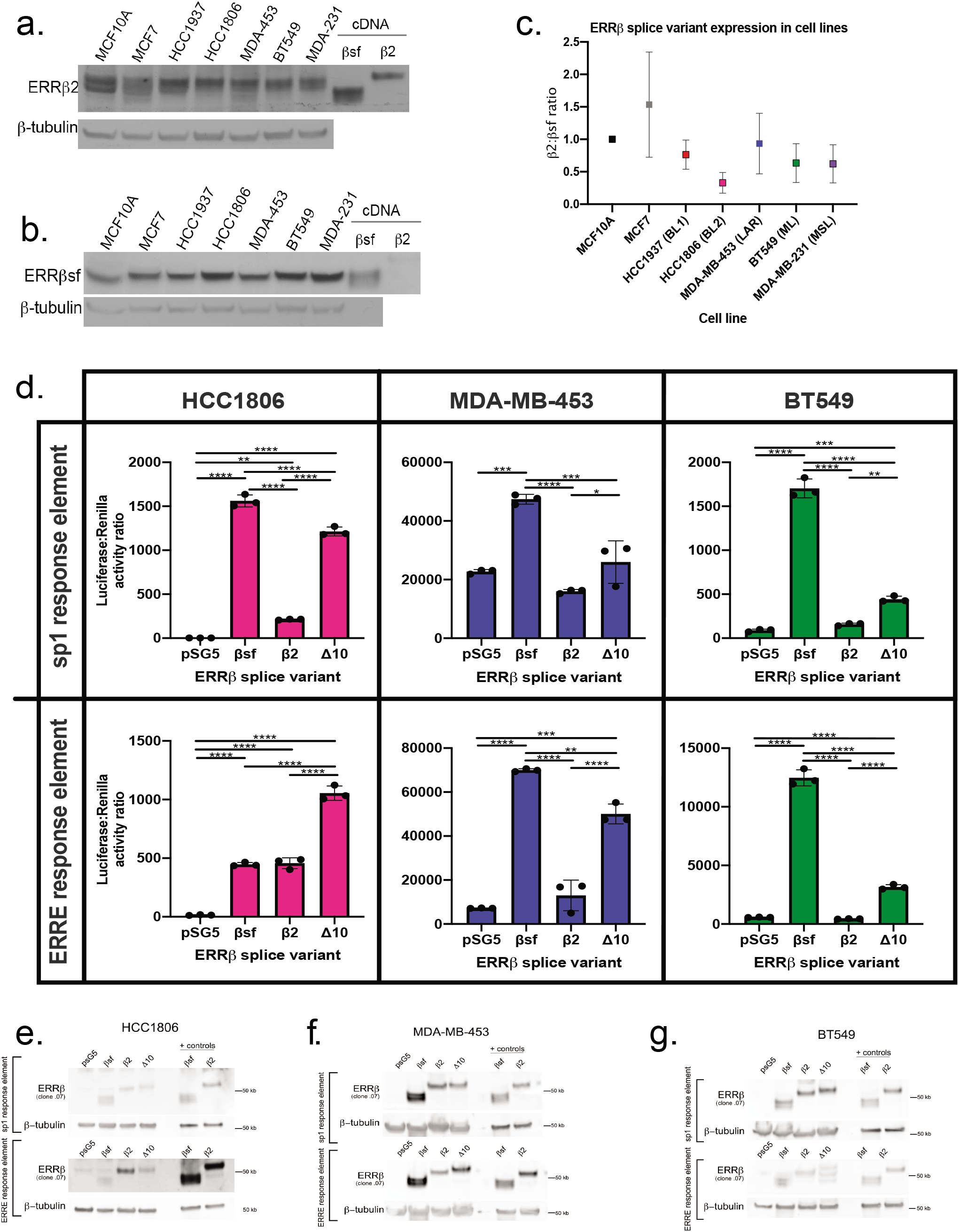
ERRβ transcription factor activity measured by heterologous promoter-reporter assays. **a,b.** Endogenous expression of ERRβ2 **(a)** and ERRβsf **(b)** protein levels in cell lines representing TNBC subtypes. **c.** Ratio of ERRβ2 and ERRβsf expression in cell lines, determined by densitometry for each receptor and corrected by expression of the β-actin loading control. **(d)** Measure of luciferase activity on sp1 and ERRE response elements in TNBC cell lines HCC1806, MDA-MB-453, and BT549. Kruskal-Wallis one-way ANOVA, ** p <0.001, ***p<0.0001. **(e-g)** Western blot confirming overexpression of ERRβ splice variant plasmid DNA in cell lines.

To measure transcription factor function of ERRβ isoforms, we overexpressed cDNAs for each isoform in three different TNBC cell lines and measured their ability to induce transcription from heterologous promoter-reporter constructs. The three cell lines used were HCC1806, MDA-MB-453, and BT549, representing 3 of 6 TNBC subtypes [34]. The promoter-reporter constructs contained sequences corresponding to 2 established ERRβ response elements: estrogen related response element (ERRE, <u>TNAAGGTCA</u>) [36, 37] and specificity-protein-1 (SP1, (G/T)GGGCGG(G/A)(G/A)(C/T)) [38] (Figure 5d). SP1 was also one of the top motifs identified in the BLBC subtype from the SCAN-B data set (Table 1). In HCC1806 (BL2) cells, there was significantly higher activity of both the ERRE and SP1 response elements induced by all three splice variants in comparison to empty vector. ERRβsf had significantly higher activity on the ERRE than ERRβ2 or ERRβ-Δ10, while ERRβ-Δ10 had significantly higher activity on the SP1 response element than ERRβsf or ERRβ2. In MDA-MB-453 (LAR) cells, ERRβsf had significantly higher activity on both the ERRE and SP1 response elements, while ERRβ-Δ10 had significantly higher activity on SP1 in comparison to empty vector. In BT549 (mesenchymal-like, ML) cells, ERRβsf and ERRβ-Δ10 had significantly higher activity on both the ERRE and SP1 response elements in comparison to empty vector. For both MDA-MB-453s and BT549s, ERRβ2 had no significant transcription factor activity on either ERRE or SP1. Western blot analysis confirmed overexpression of ERRβ isoforms in transfected cells (Fig. 5e-g). These data suggest that patterns of ERRβ isoform transcription factor activity may differ between basal-like and other TNBC molecular subtypes.

### ERRβ isoform expression in breast tumor tissue varies by IHC subtype

We next assessed the protein expression of ERRβsf and ERRβ2 isoforms in patient tissues from a 150-patient TMA assembled by the HTSR at Georgetown University Medical Center’s LCCC. The TMA series consists of 50 invasive ductal carcinoma breast cancer patients each diagnosed with ER+, HER2 overexpressing, and TNBC IHC subtypes, with corresponding demographic and clinical information (race, pathological stage, age, treatment, Table 2). Post antibody optimization, consecutive sections of the TMA were stained for 1 of 2 antibodies: ERRβ-clone 05 (recognizes ERRβsf) or ERRβ-clone 07 (recognizes ERRβ2). Staining was quantified in a semiautomated fashion and nuclear versus cytoplasmic staining were differentiated using the Vectra3 Multi-Spectral Imaging Microscope. (Fig. 6a-c). Representative image show the differential staining of the two antibodies (Fig. 6d). Statistical analyses were performed to determine the relationship of each isoform with lymph node status, race, and age.

**Table 2.**
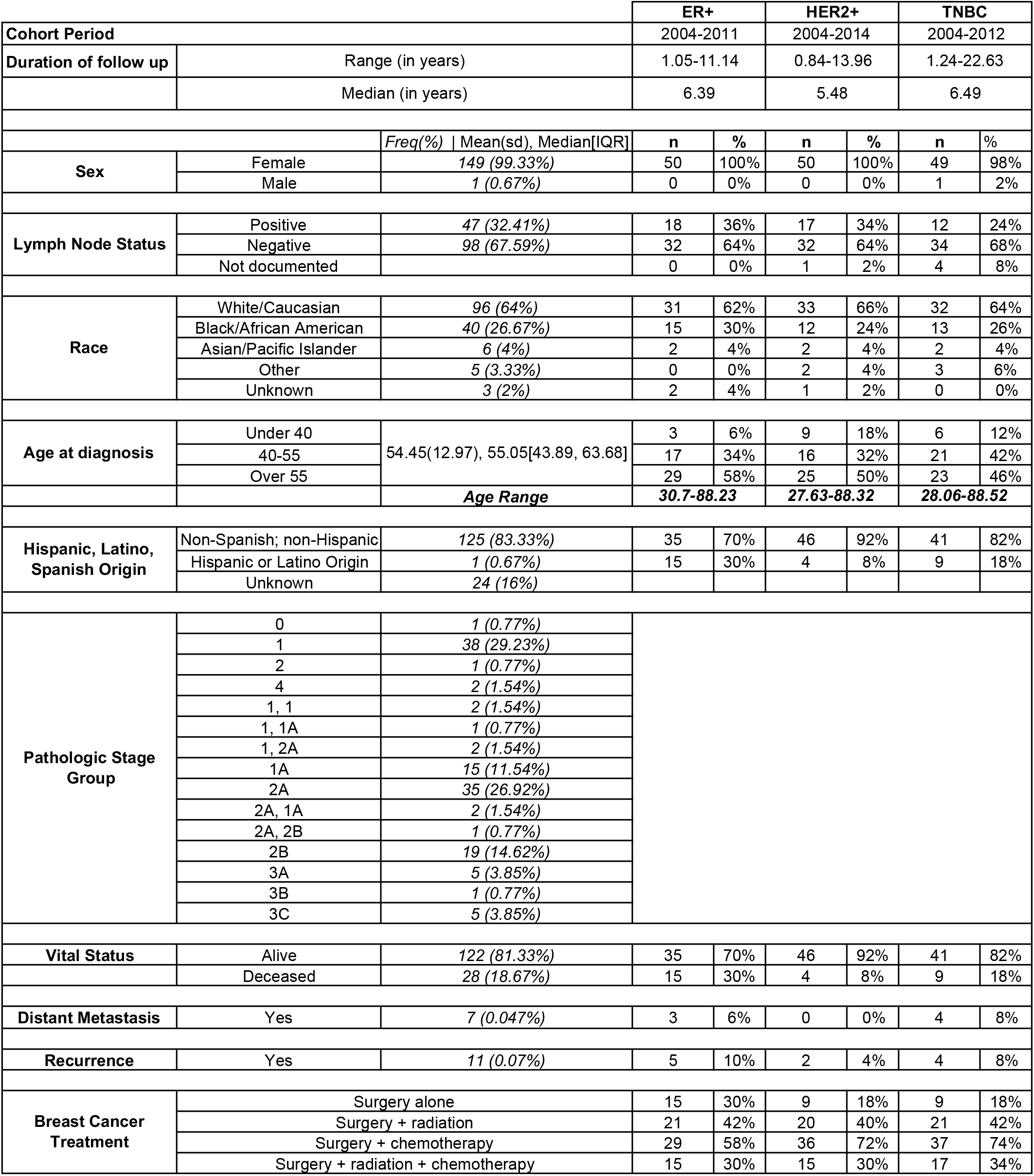
Demographics of the 150-patient cohort included in the invasive ductal carcinoma (IDC) tissue microarray (TMA).

**Fig. 6.**
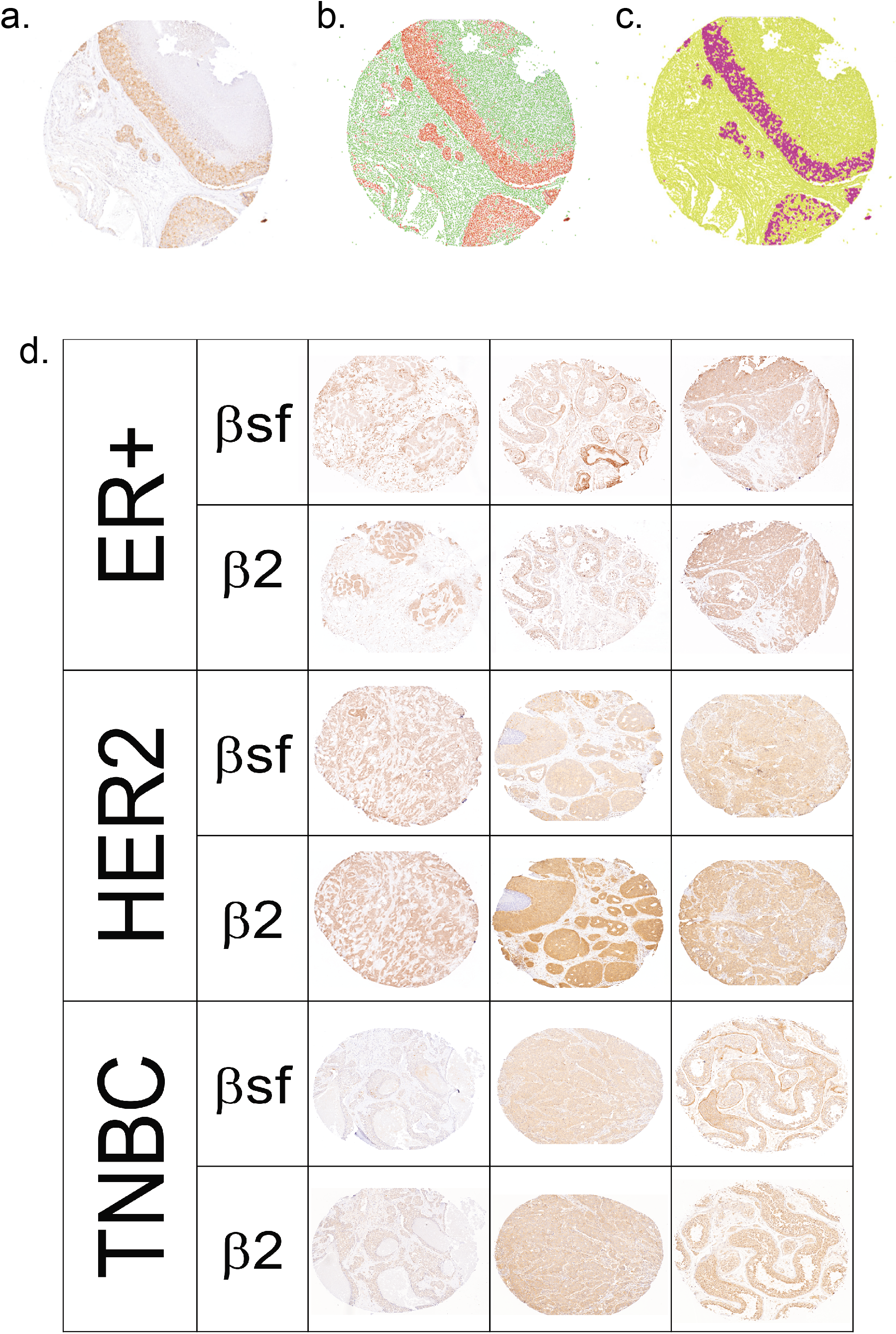
ERRβ isoform expression by IHC. A-D. Automatic scanning and semiautomatic quantification of ERRβ splice variant specific mouse monoclonal antibodies (ab) on tissue microarray (TMA). **a.** Blue/brown staining with ERRβ ab. Automatic detection of **b.** nuclear v cytoplasmic staining, and **c.** cells ‘‘positive”/ above threshold vs. “negative”/ below threshold. **d.** ERRβ protein expression in breast tumors. Representative images of ER+, HER2, and TNBC tumor tissues from 150 patient tissue microarray series stained with ERRβ ab

The median of ERRβ2, but not ERRβsf, expression was significantly different amongst the three IHC receptor subtypes: ER+, HER2, TNBC (p = 0.016). Post hoc analysis showed that ERRβ2 was significantly different between ER+ and HER2 patients (p = 0.017, Figure 7a, Table S2.1). Median ERRβsf: ERRβ2 expression ratio was lowest in TNBC subtypes (indicating lower ERRβ2 or higher ERRβsf expression), however this ratio was not statistically significantly different amongst the three IHC receptor subtypes (Figure 7b).

**Fig. 7.**
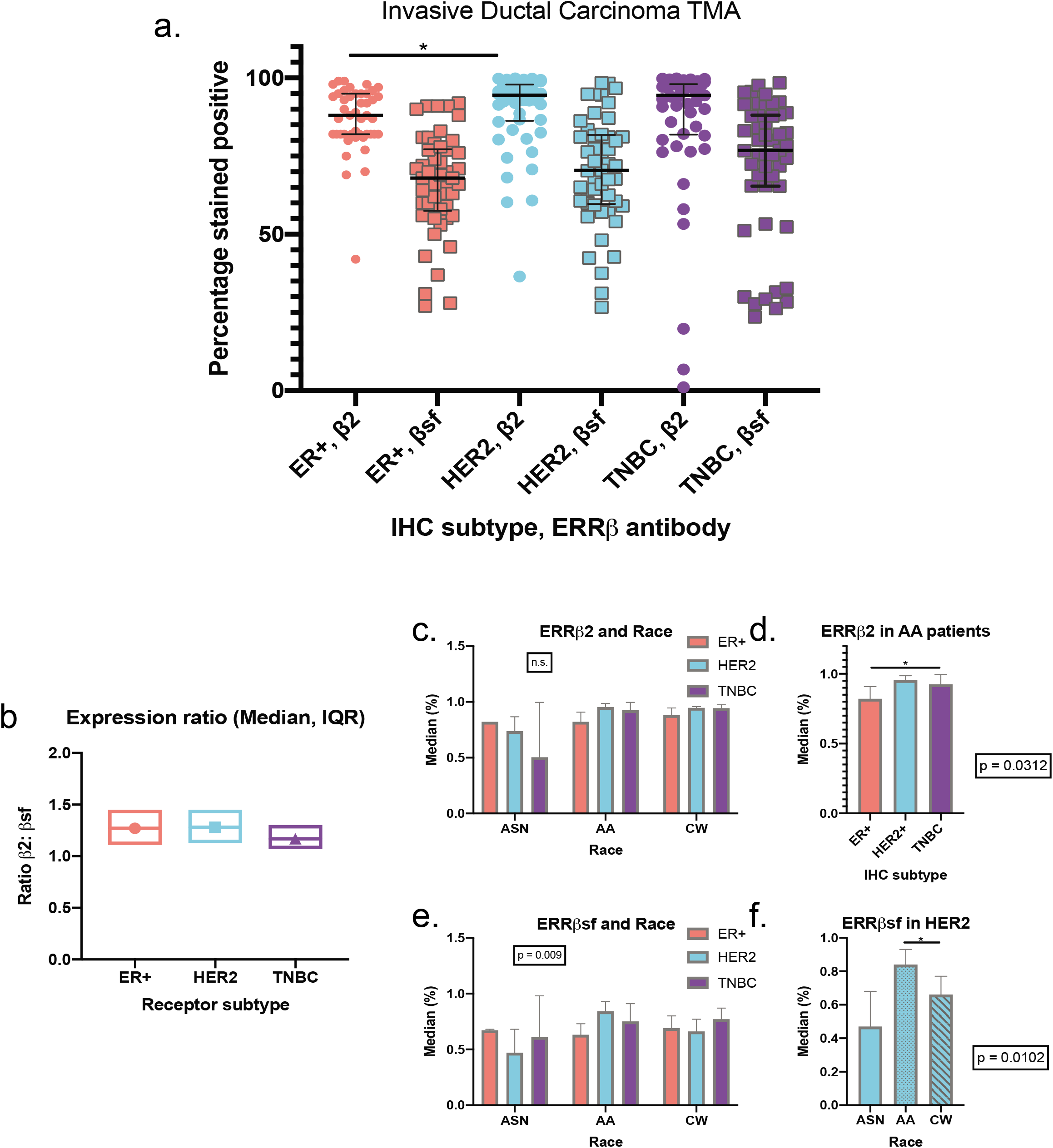
Semi-automated quantification of ERRβ IHC. **A.** Total percent (%) of patients that are ERRβ positive, by IHC subtype, *p<0.01. **b.** Ratio of total ERRβ2:total ERRβsf determined for ER+, HER2+, and TNBC patients. **c-e.** ERRβ expression by race. Interactions between ERRβ2 receptor expression **(c,d)** and ERRβsf expression **(e,f)** and race. Two-way ANOVA, followed by DSCF pair-wise comparison *p < 0.05

We next assessed statistical interactions and main effects of three variables: IHC subtypes (variable 1), lymph node status, race or age (variable 2), and ERRβ status (variable 3). For ERRβ2 expression, there were no statistical interactions between the IHC subtypes and lymph node status, race or age that affected ERRβ status. There was no significant main effect on ERRβ2 expression by IHC subtypes with lymph node status, or, by IHC subtypes with age. However, race did show a significant main effect (p = 0.024) or difference in ERRβ2 expression amongst the different races (Fig. 7c, Table S3). Follow-up post-hoc Kruskal-Wallis found that the ERRβ2 median was significantly different amongst the three IHC subtypes in AA patients (p = 0.0312, Figure 7d). Further pairwise comparisons with DSCF method showed this observed difference was between ER+ and TNBC patients (p = 0.0212, Figure 7b, d).

There were no significant statistical interactions between the IHC subtypes and lymph node status or age that affected ERRβsf status, however we did see significant interaction effect with race (p = 0.009) indicating differential ERRβsf expression within the 5 races between the 3 IHC subtypes (Table S3, Fig. 7e). This significant interaction was followed up with a Kruskal-Wallis post hoc analysis which showed that median ERRβsf expression was significantly different among 5 races in the HER2 IHC receptor subtype (p = 0.0102). Follow-up pairwise comparisons showed that this significant difference was between AA and CW patients (p = 0.0282, Figure 7f).

We next analyzed the subcellular localization of ERRβsf and ERRβ2 isoforms amongst the different IHC subtypes. Previous studies have shown that ERRβsf and ERRβ2 have different localization and functions within cells [11], with ERRβsf predominantly nuclear and ERRβ2 expressed in both the nucleus and cytoplasm. ERRβ2 had higher cytoplasmic vs. nuclear staining in ER+ patients compared to HER2 and TNBC patients, while ERRβsf had both nuclear and cytoplasmic staining in all three IHC subtypes (Fig. 8a, 8b, Table S2.2). We performed Spearman rank analysis to measure the association between the nuclear and cytoplasmic staining of ERRβ2 and ERRβsf in each IHC subtype. In the ER+ group, both ERRβ2 and ERRβsf nuclear staining were significantly positively correlated with cytoplasmic staining (r= 0.945, p < 0. 0001 and r=0.488, p = 0.00038, respectively) meaning the expression of nuclear ERRβ2/ERRβsf increases when the value of cytoplasmic ERRβ2/ERRβsf increases. In the HER2 group, ERRβ2 nuclear staining was significantly negatively correlated with cytoplasmic staining (r= −0.922, p <0.0001), while ERRβsf nuclear staining was positively correlated with cytoplasmic staining (r=0.928, p <0.0001). In the TNBC group, ERRβ2 nuclear staining was also significantly negatively correlated with cytoplasmic staining (r= −0.729, p <0.0001), while ERRβsf nuclear staining was positively correlated with cytoplasmic staining, (r=0.93, p <0.0001). (Figure 8c).

**Fig. 8.**
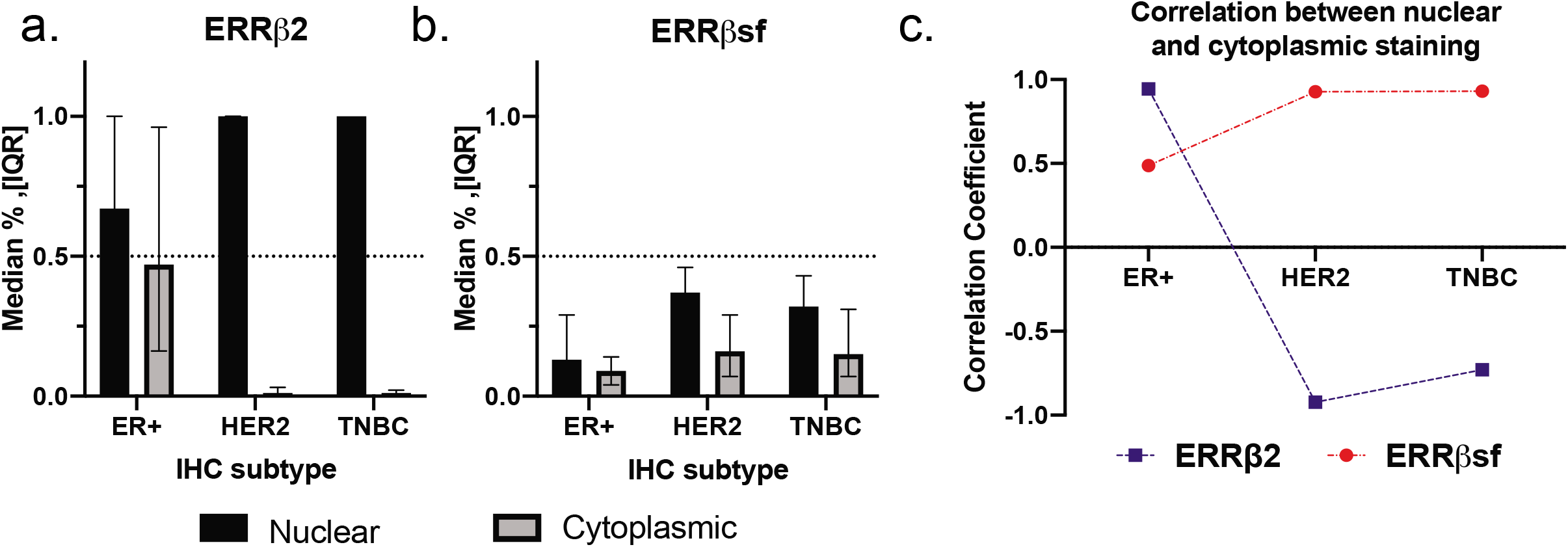
Subcellular localization of ERRβ2 and ERRβsf in TMA cores. Cytoplasmic and nuclear localization of ERRβ2 **(a)** and ERRβsf **(b)** determined for the three IHC subtypes. **c.** Correlation of ERRβ2 and ERRβsf between nuclear and cytoplasmic staining, by IHC subtype

## DISCUSSION

In this study, we characterize ONR ESRRB/ERRβ copy number and expression and its clinical correlations in breast cancer. Patients with low *ESRRB* mRNA expression have significantly shorter overall survival. *ESRRB* mRNA expression is significantly lower in TNBC/BLBC vs. other breast cancer subtypes, but not significantly associated with age, lymph node status, or grade. In an independent dataset, we find no evidence of *ESRRB* copy number loss in TNBC patients, suggesting that reduced mRNA expression is driven by other mechanisms. We further find that *ESRRB* expression is correlated with that of genes associated with neuroactive ligand-receptor interaction, metabolic pathways, and deafness, and that these genes contain G/C-rich transcription factor binding motifs. We show that the *ESRRB* message, which is alternatively spliced into three distinct isoforms, leads to different isoform transcription factor activity in TNBC cell lines that are characterized as basal-like vs. the mesenchymal or luminal androgen receptor subtype. Finally, we show in clinical samples that the ERRβ2 and ERRβsf isoforms are broadly expressed at the protein level.

We found that the loss of *ESRRB* mRNA in TNBC/BLBC patients was not due to a loss of copy number, and in fact there was a gain in *ESRRB* copy number in all breast cancer subtypes. Though the copy number gain was not significant, it does suggest that there is a mechanistic variance occurring during transcription of this gene in breast cancer patients that drives the loss of mRNA that is consistently and significantly observed especially in TNBC/BLBC. While our ancestrality marker results were not significant, in the subset, which had a small number of patients, we saw a trend toward reduced *ESRRB* copy number in AA versus CW patients which aligned with 1000 genome projects defined AFR and AMR patients, respectively. This addresses a clinical diagnosis issue frequently encountered in which demographic information is misreported [39]. In this case, we looked at self-reported race in comparison to refined race and ethnic information from genotyping, and saw a difference in copy number that was not seen with only the self-reported race. Our results suggest that there is value to the addition of ancestrality markers to future analyses would add more insight.

When looking at transcription factor activity we found that the three known splice variants ERRβsf, ERRβ2, ERRβ-Δ10 when exogenously expressed, have differential activity on known *ESRRB* response elements. In basal-like TNBC cell line, BL2, ERRβsf had higher activity on the ERRE while ERRβ-Δ10 had higher activity on the sp1 response element. Meanwhile in LAR and ML TNBC cell lines, ERRβsf and ERRβ-Δ10 had higher activity on both response elements, while ERRβ2 had no change in activity. The LAR and ML subtypes are consistent with our previous publication in which we showed that in a ER+ setting ERRβsf has higher activity on the ERRE while ERRβ2 shows no change in activity [11]. The difference in activity may be explained by the differing C-terminus of the splice variants. While all three isoforms share exons 3 - 9, ERRβsf is truncated at exon 9, ERRβ2 has exon 10 and part of 11 and ERRβ-Δ10 skips exon 10, but contains exons 11 and 12. Due to the different exon combinations, the 3 variants have different functional domains which may cause steric hindrance and thus may alter the binding of each variant to DNA. To further validate these findings, stable cell lines with inducible overexpression of *ESRRB* isoforms should be established. Transient overexpression limits the time frame in which experiments can be carried out to 24-72 hours, with optimal expression ~48 hours in the TNBC cell lines. Stable overexpression would allow us to further evaluate other features such as proliferation, cell death, migration, and invasion in association with overexpression of specific *ESRRB* isoforms. Non-transformed mammary epithelial cells MCF10As should also be established with both stable overexpression and knockdown/knockout with rescue.

This is the first reported comparative quantification of ERRβsf and ERRβ2 protein expression in IHC breast cancer subtypes. Our 150-patient TMA revealed that in all three IHC subtypes (ER+, HER2, TNBC) ERRβ2 is expressed in higher quantities than ERRβsf. We established that ERRβ2 was significantly different between ER+ and HER2 patients, and that the ratio of ERRβ2:ERRβsf was lower in TNBC patients that it was in ER+ or HER2 patients. Though this difference was not statistically significant, it did suggest that in the more aggressive breast cancer subtype there is an increase in the ERRβsf splice variant. Our previous publications have led to the hypothesis that ERRβ specific - ligand-mediated mitotic arrest is due to the ERRβ2 splice variant, and that the loss of this splice variant is an effect of the malignant transformation associated with TNBC/BLBC [11].

We have previously published that in breast cancer cell lines ERRβ2 localizes to the cytoplasm and centrosomes in the nucleus while ERRβsf localizes in the nucleus [11]. We further looked at this localization to see if there were differences amongst the three IHC subtypes. We established that while in ER+ tumors there was a positive correlation between nuclear and cytoplasmic staining for ERRβ2 and ERRβsf, both HER2 and TNBC - clinically aggressive subtypes - showed a positive correlation for ERRβsf and a negative correlation for ERRβ2. Furthermore, ERRβ2 localized almost exclusively in the nucleus in both of those settings. The inconsistency from our previous findings points to the question of heterogeneity and how to best treat cancer patients based on the different cells within a tumor.

This study has provided insight into the expression of ERRβ and its implications in TNBC/BLBC, an aggressive subtype of breast cancer which, to date, does not have any targeted therapies. Limitations of our study include the underrepresentation of African-American patients, a group that is disproportionally affected by this breast cancer subtype. Though the SCAN-B data set is very large, it has limited racial and ethnic diversity. Our TMA had ~1/3 AA patients, an improvement over past studies some of which do not include any AA patients. Additionally, we have shown the value ancestrality information can provide in lieu of self-reported race.

Another limitation of our analyses includes the inability to properly examine splice variant – specific *ESRRB* at the mRNA level. Due to the overall loss of *ESRRB* observed in the cancer setting, it is extremely difficult to look at individual isoform expression. To properly quantify splice variants at the RNA level, we would need to use ultra-deep RNAseq [40, 41]. This increases genome coverage and improves sequencing accuracy, providing assurance that we are accurately looking at isoform mRNA expression.

Future steps include *in vitro* validation of our DEG, pathway, and promoter analyses to further determine *ESRRB* co-regulatory partners. As previously stated, the establishment of stable cell lines will be instrumental in these validation studies. With this information ESRRB/ERRβ could serve as a future therapeutic target or as a biomarker in TNBC.

## Acknowledgments

We are grateful to Allison O’Connell, Dr. Hillary Stires, Ayodeji Olukoya, and Sonali Persaud for their insights and/or critical reading of the manuscript. We would like to thank Drs. Bassem Haddad, Filipa Lynce, and Michael Johnson for their guidance in developing the HTSR’s invasive ductal carcinoma breast cancer tissue microarray series. We thank Henry Cho and Gaelle Palmer for their contributions to the TMA-associated REDCap database. Thank you to the Survey, Recruitment, and Biospecimen collection Shared Resource (SRBSR) for their support of research recruitment at MedStar Georgetown University Hospital. Thank you to Dr. Lao Saal of Lund University, Sweden for kindly providing the clinical information of SCAN-B data. We thank Garrett Graham for his guidance on all computational studies and Max Kushner for his aid with the DREME analysis. The results shown here are based in part upon data generated by the TCGA Research Network: https://www.cancer.gov/tcga.

## ABBREVIATIONS

*TNBC*: Triple negative breast cancer
*BLBC*: Basal-like breast cancer
*AA*: African-American
*ESRRB*: Estrogen related receptor beta
*IHC*: Immunohistochemistry
*ER*: Estrogen receptor
*PR*: progesterone receptor
*HER2*: human epidermal growth factor two
*CW*: Caucasian/White
*NR*: Nuclear receptor(s)
*ONR*: Orphan nuclear receptor
*ERR*: Estrogen related receptors
*OS*: Overall survival
*SCAN-B*: Sweden Cancerome Analysis Network - Breast
*FPKM*: Fragments Per Kilobase of transcript per Million mapped reads
*ESR1*: Estrogen receptor
*NHG*: Nottingham grade
*NTN*: Non triple negative breast cancer
*aCGH*: Array comparative genomic hybridization
*AFR*: African descent
*AMR*: Ad mixed American
*DEGs*: Differentially expressed genes
*BL2*: Basal-like 2
*LAR*: Luminal Androgen Receptor
*ML*: Mesenchymal-like
*ERRE*: Estrogen related response element
*SP1*: Specificity-protein-1
*TMA*: Tissue microarray

