## Supplementary Figure and Table Legends for "The orphan nuclear receptor estrogen-related receptor beta (ERRβ) in triple-negative breast cancer"

**Supplemental Fig. 1. Age at diagnosis and association of *ESRRB* with OS in SCAN-B and TCGA data sets**. **a, b.** Average age of patients in SCAN-B data set by Pam50 **(a)** and IHC **(b)** subtypes. ANOVA with multiple comparisons ***p<0.001, **** p<0.0001. **c-h.** KM plots showing overall survival of all breast cancer patients **(c,f)**, BLBC patients **(d,g)**, and TNBC patients **(e,h)** in SCAN-B (c-e) and TCGA (**f-h**) data

**Supplemental Fig. 2. Demographics of aCGH cohort.** **A-E.** Distribution of (**a)** age, **(b)** tumor size and *ESRRB* copy number, **(c)** pathology, **(d)** lymph node status and **(e)** metastasis

**Supplemental Table 1. DEGs in *ESRRB* high and low patients.** List of DEGs found in BLBC and TNBC patients from SCAN-B and TCGA data sets.

**Supplemental Fig. 3. Overlap of BLBC patients and TNBC patients. a, b.** Heat map representing differential expression of overlapping genes in SCAN-B versus TCGA data sets. **c, d.** Venn diagram of patients in SCAN-B **(c)** and TCGA **(d)** data. **e.** List of overrepresented motifs in promoter region DEGs in SCAN-B data sets

**Supplemental Fig. 4. Protein and mRNA levels of ERRβ / *ESRRB*.** Densitometry quantifying ERRβ splice variants ERRβ2 **(a)** and ERRβsf **(b)** protein levels in cell lines **c.** *ESRRB* mRNA levels in SCAN-B patients, sorted into TNBC subtypes. **d, e.** *ESRRB* mRNA levels in cell lines representing the TNBC cell lines

**Supplemental Table 2. ERRβ isoform expression and subcellular localization.** Mean (sd) and median (IQR) expression of ERRβ2 receptor, ERRβsf receptor, and total ERRβ2:total ERRβsf expression (S2.1) and nuclear/ cytoplasmic localization of ERRβ2 receptor and ERRβsf (S2.2) in three IHC breast cancer subtypes

**Supplemental Table 3. ERRβ isoform expression and clinical features.** Analysis of ERRβ2 and ERRβsf receptor expression in the IHC subtypes and lymph node status, race, and age, with and without interaction
