## Supplementary Figures and Tables for "The orphan nuclear receptor estrogen-related receptor beta (ERRβ) in triple-negative breast cancer"

Supplemental Fig. 1

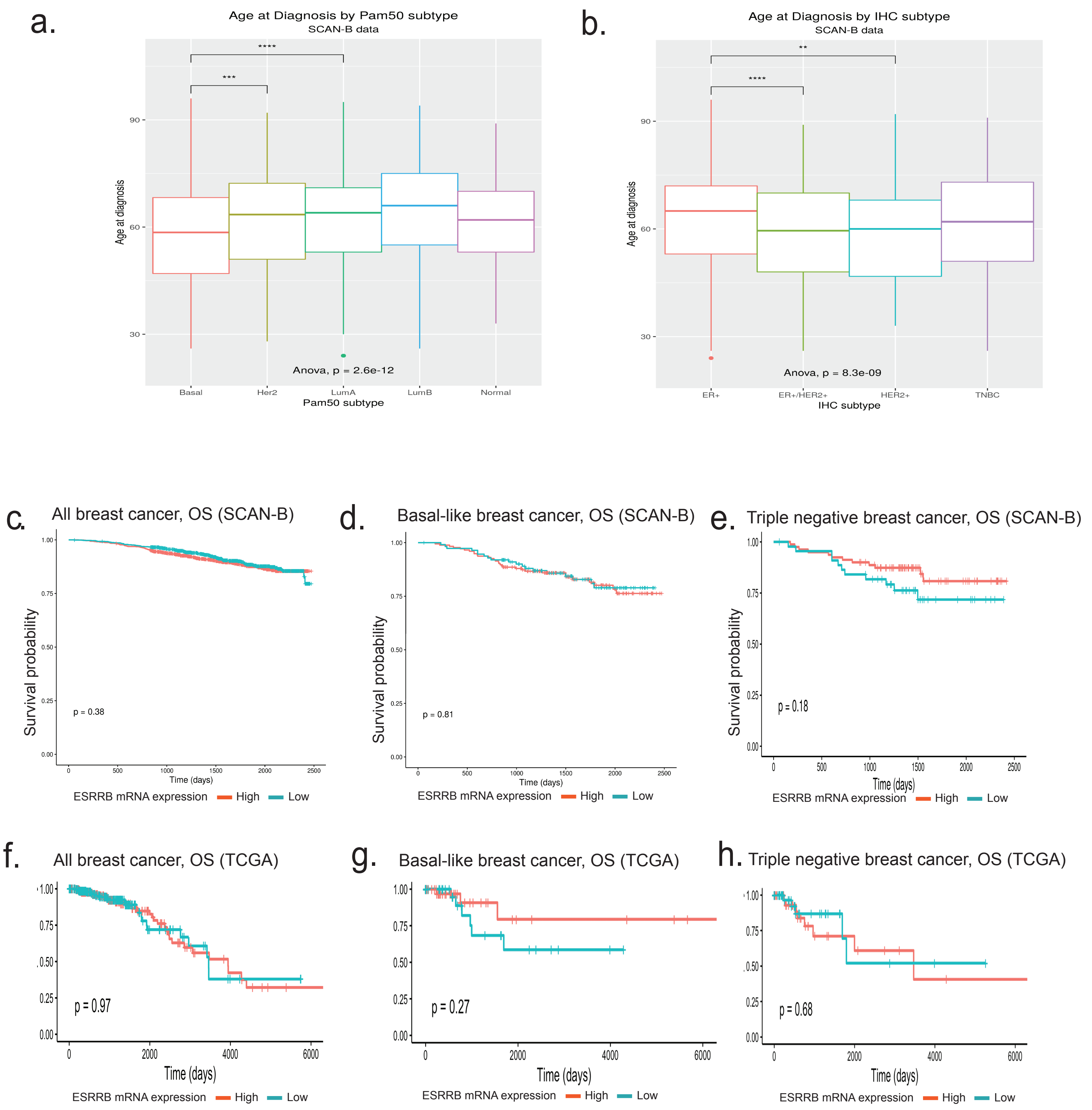

Supplemental Fig. 2

a.

### Age

| Race | mean | Classification | n | mean |
|---|---|---|---|---|
| **CW** | 53.68 | **TNBC** | 17 | 56.76 |
| | | **NTN** | 20 | 52.59 |
| **AA** | 53.23 | **TNBC** | 39 | 53.29 |
| | | **NTN** | 30 | 54.18 |

b.

### Mean *ESRRB* CN by Tumor size

| | | **CW, mean=0.775** | | | | | | **AA, mean=0.591** | | | | | |
|---|---|---|---|---|---|---|---|---|---|---|---|---|---|
| | | TNBC | | | NTN | | | TNBC | | | NTN | | |
| **Tumor size** | **Correlative stage** | **mean** | **sd** | **n** | **mean** | **sd** | **n** | **mean** | **sd** | **n** | **mean** | **sd** | **n** |
| >0.1 ≤ 0.5 | T1a | - | - | - | 1.197 | 0.000 | 1 | 0.580 | 0.000 | 1 | 0.162 | 0.000 | 1 |
| > 0.5 ≤ 1 | T1b | 0.829 | 0.000 | 1 | 1.330 | 0.451 | 2 | 0.760 | 0.247 | 10 | 0.945 | 0.452 | 9 |
| > 1 ≤ 2 | T1c | 0.761 | 0.906 | 6 | 0.630 | 0.681 | 11 | 0.459 | 0.612 | 14 | 0.731 | 0.848 | 12 |
| > 2 ≤ 5 | T2 | 0.224 | 0.262 | 5 | 0.943 | 0.491 | 6 | 0.165 | 0.732 | 10 | 0.547 | 0.418 | 5 |
| >5 | T3 | 0.285 | 0.112 | 3 | - | - | - | 0.882 | 0.267 | 3 | 0.676 | 0.365 | 2 |

c.

### Pathology

| Race | Classification | Pathology | % patients |
|---|---|---|---|
| **CW** | **TNBC** | In situ/IDC | 23.5% |
| | | IDC | 64.7% |
| | | Other | 11.8% |
| | **NTN** | In situ/IDC | - |
| | | IDC | 80.0% |
| | | Other | 20.0% |
| **AA** | **TNBC** | In situ/IDC | 30.8% |
| | | IDC | 69.2% |
| | | Other | - |
| | **NTN** | In situ/IDC | 60.0% |
| | | IDC | 23.3% |
| | | Other | 16.7% |

d.

### Lymph node status

| | | **% patients** | | |
|---|---|---|---|---|
| **Race** | **Classification** | **Negative** | **Positive** | **NA** |
| **CW** | **TNBC** | 47% | 29% | 24% |
| | **NTN** | 45% | 40% | 15% |
| **AA** | **TNBC** | 56% | 44% | 0% |
| | **NTN** | 50% | 37% | 13% |

e.

### Metastasis

| | | **% patients** | | |
|---|---|---|---|---|
| **Race** | **Classification** | **None** | **Local** | **Distant** |
| **CW** | **TNBC** | 59% | 24% | 18% |
| | **NTN** | 45% | 30% | 25% |
| **AA** | **TNBC** | 62% | 21% | 18% |
| | **NTN** | 63% | 23% | 13% |

Supplemental Table 1

| SCAN-B, BLBC | | |
|---|---|---|
| Gene | logFC | P.Value |
| SPTBN5 | 7.639370222 | 0.000873051 |
| FLG | 6.994851245 | 0.030036705 |
| BC065766 | 6.718097379 | 0.031511173 |
| SFTPA2 | 6.27596873 | 0.00234021 |
| TMEFF2 | 5.935253136 | 0.029022906 |
| CIB4 | 5.542421462 | 0.006344489 |
| TLL2 | 5.502385296 | 0.006993371 |
| IFITM5 | 5.239709767 | 0.041785865 |
| TIPARP | 5.021939147 | 0.039978781 |
| SLC8A1 | 5.018940109 | 0.043888801 |
| AK091729 | 4.779418674 | 0.046404664 |
| NTN3 | 4.699663842 | 0.006911973 |
| FAM9C | 4.697800912 | 0.007437151 |
| NXNL2 | 4.673342092 | 0.019596239 |
| LOC100506730 | 4.652089627 | 0.047854364 |
| AK094577 | 4.593986107 | 0.000103129 |
| TBX10 | 4.525318833 | 0.013276925 |
| TMIE | 4.517565838 | 0.029484905 |
| GRIK1 | 4.47475979 | 0.02821595 |
| PRKAG3 | 4.331722869 | 0.026115227 |
| SLC6A4 | 4.124948641 | 0.04062148 |
| LPAR4 | 4.056102105 | 0.04268291 |
| ANXA13 | 4.052885868 | 0.011945107 |
| FGB | 4.025989727 | 0.005187143 |
| MAR4 | 3.870006874 | 0.043287681 |
| AK024736 | 3.774088451 | 0.014528722 |
| AK128708 | 3.692015861 | 0.001341628 |
| GRIA4 | 3.651999261 | 0.02865887 |
| LINC00491 | 3.603976673 | 0.036705444 |
| DKFZp434J0226 | 3.558614319 | 0.010748635 |
| MYOM2 | 3.540636408 | 0.025883866 |
| AK124970 | 3.358091144 | 0.049041097 |
| LHX1 | 3.350989863 | 0.027099116 |
| LINC00964 | 3.323078307 | 0.006463981 |
| LOC389705 | 3.308892993 | 0.01250708 |
| NKX2 | 3.212344494 | 0.006488279 |
| BC033241 | 3.145756283 | 0.033993858 |
| AK021876 | 3.135691395 | 0.052528899 |
| SLC5A11 | 3.096699035 | 0.010702889 |
| AK097921 | 3.045190181 | 0.003364293 |
| BC044608 | 3.03622691 | 0.012132061 |
| LOC155060 | 2.826289683 | 0.014577276 |
| CT45A1 | 2.820927706 | 0.001908482 |
| LINC00886 | 2.789380613 | 0.003908043 |
| AK128252 | 2.785083511 | 0.022883708 |
| ANKRD30B | 2.690610855 | 0.037723802 |
| NCR1 | 2.682998696 | 0.002128345 |
| GAST | 2.675539776 | 0.020999989 |
| CETN4P | 2.666792752 | 0.037100539 |
| LOC101059948 | 2.651440569 | 0.001045381 |
| SIGLEC17P | 2.636115466 | 0.029118709 |
| LRRC46 | 2.562091538 | 0.026033824 |
| BC141952 | 2.508584977 | 0.026633284 |
| KIAA2022 | 2.499643714 | 0.013209174 |
| KF274612 | 2.481023208 | 0.023280248 |
| AL832163 | 2.466024503 | 0.000275446 |
| KCNH3 | 2.445352169 | 0.007857042 |
| ABHD1 | 2.42579731 | 0.050058955 |
| CBLN2 | 2.383375531 | 0.030188307 |
| BC068290 | 2.365398509 | 0.025128632 |
| TAT | 2.344279688 | 0.002168722 |
| OTOF | 2.342409809 | 0.001042041 |
| AK311558 | 2.341525373 | 0.019807594 |
| RGS9BP | 2.264883564 | 0.04496302 |
| DQ593432 | 2.246610581 | 0.041841021 |
| CYP4F22 | 2.217945918 | 0.010289675 |
| VSIG8 | 2.186892825 | 0.022232452 |
| PACRG | 2.150946354 | 0.004838196 |
| FCRL1 | 2.097513229 | 0.002802256 |
| MIR29C | 2.073467429 | 0.044759358 |
| KCNE2 | 2.022137329 | 0.008502552 |
| EGOT | 2.010978885 | 0.000112788 |
| SFTA1P | -2.075774235 | 0.035199114 |
| AK092862 | -2.092409032 | 0.03439314 |
| BC069004 | -2.095062307 | 0.009578383 |
| LCE3D | -2.106312802 | 0.028925308 |
| CNTNAP3B | -2.119882001 | 0.015782902 |
| NPBWR1 | -2.128822225 | 0.022033416 |
| MS4A8 | -2.129663717 | 0.006067197 |
| RP11 | -2.154236682 | 0.018675995 |
| PARD3 | -2.205534591 | 0.01748693 |
| SPRR4 | -2.231220213 | 0.027253698 |
| FIGLA | -2.271935823 | 0.031572529 |
| FSTL5 | -2.305181207 | 0.034505526 |
| PTPRR | -2.309334031 | 0.031935166 |
| PHYHIP | -2.321287084 | 0.052782296 |
| TAPT1 | -2.323169452 | 0.018752984 |
| KIAA0319 | -2.334514256 | 0.022963902 |
| C6orf165 | -2.357103563 | 0.047830985 |
| PCDHGA1 | -2.391610048 | 0.010382972 |
| CFTR | -2.400518371 | 0.035090334 |

| SCAN-B, TNBC | | |
|---|---|---|
| Gene | logFC | P.Value |
| SPTBN5 | 8.474119915 | 0.00276 |
| EXOC3L4 | 8.03463058 | 0.00008 |
| KIAA2022 | 5.909117267 | 0.00019 |
| UGT2B15 | 5.198585536 | 0.04513 |
| PAH | 4.903177801 | 0.00259 |
| IL17REL | 4.801767819 | 0.00336 |
| TRIML2 | 4.668769472 | 0.01595 |
| RP11 | 4.502733507 | 0.03916 |
| TBX10 | 4.472770595 | 0.00959 |
| AK093002 | 4.428159445 | 0.01006 |
| LOC494141 | 3.979636105 | 0.03110 |
| GRIK5 | 3.943006643 | 0.04724 |
| ATP10B | 3.901132671 | 0.02228 |
| FGB | 3.863986331 | 0.01190 |
| AK056887 | 3.719072657 | 0.00493 |
| ABHD1 | 3.713649521 | 0.03670 |
| GRIN3A | 3.706993954 | 0.02539 |
| DEFB124 | 3.543091681 | 0.03805 |
| ZP1 | 3.522836667 | 0.01918 |
| AK123450 | 3.516912447 | 0.00428 |
| CLEC9A | 3.513366355 | 0.02036 |
| TPO | 3.497630719 | 0.01047 |
| GLOD5 | 3.426685794 | 0.02329 |
| RASA4CP | 3.348876404 | 0.01468 |
| RPH3A | 3.245058593 | 0.00579 |
| GUCA1B | 3.232226545 | 0.02331 |
| C19orf83 | 3.191135457 | 0.01322 |
| SLC6A10P | 3.145316915 | 0.00038 |
| FCRL1 | 2.997812435 | 0.03861 |
| AQP4 | 2.99580351 | 0.00601 |
| LOC155060 | 2.991451077 | 0.05002 |
| TAT | 2.913089419 | 0.01823 |
| LRRC46 | 2.84854943 | 0.01623 |
| CSN1S1 | 2.800142359 | 0.01606 |
| SCGB1A1 | 2.749999611 | 0.04515 |
| LINC00202 | 2.747357412 | 0.02140 |
| CFHR1 | 2.71708801 | 0.04710 |
| SEMA3A | 2.702752905 | 0.00592 |
| KRT9 | 2.637140327 | 0.01601 |
| AL592528 | 2.631859891 | 0.01351 |
| BC036382 | 2.630416346 | 0.03850 |
| BX648455 | 2.427905175 | 0.03460 |
| RGS9BP | 2.426216685 | 0.02888 |
| PACRG | 2.403796277 | 0.01022 |
| LOC100506022 | 2.368848311 | 0.04098 |
| RP11 | 2.343006996 | 0.00004 |
| HPSE2 | 2.341521669 | 0.03457 |
| ADAMTS8 | 2.320057907 | 0.00719 |
| PLD5 | 2.31540911 | 0.01285 |
| BCAN | 2.283520792 | 0.00526 |
| STK32B | 2.267200192 | 0.00626 |
| LOC100130298 | 2.257475943 | 0.02049 |
| RPRM | 2.254338594 | 0.03880 |
| CYP4F22 | 2.231054987 | 0.00107 |
| BC062291 | 2.217284554 | 0.01233 |
| COL23A1 | 2.214001812 | 0.00032 |
| BC040901 | 2.130167212 | 0.02284 |
| SLC25A21 | 2.122326393 | 0.00040 |
| AGR3 | 2.077650249 | 0.00286 |
| GREB1L | 2.070130776 | 0.00702 |
| PPP1R36 | -2.005576658 | 0.01689 |
| MKRN9P | -2.089792473 | 0.01668 |
| TNFSF12 | -2.096986869 | 0.00258 |
| DNASE1L2 | -2.124655199 | 0.01402 |
| IDI2 | -2.146826115 | 0.04456 |
| BC035726 | -2.201031184 | 0.00289 |
| TMEM151B | -2.205214875 | 0.04913 |
| PFKFB1 | -2.205971096 | 0.03231 |
| BC033961 | -2.211363702 | 0.03733 |
| LOC100128398 | -2.223668441 | 0.00749 |
| LOC100289333 | -2.227575745 | 0.02743 |
| FCN2 | -2.2700934 | 0.02713 |
| EML5 | -2.292035404 | 0.04741 |
| ABCA17P | -2.302978936 | 0.00079 |
| ACVR2B | -2.342205582 | 0.03647 |
| TPRG1 | -2.366380985 | 0.02611 |
| IL22RA1 | -2.369708498 | 0.03686 |
| PNLIPRP3 | -2.376175473 | 0.02572 |
| MYOM2 | -2.387244692 | 0.00513 |
| MPO | -2.392922726 | 0.01483 |
| AKAP3 | -2.438775309 | 0.02834 |
| AK127124 | -2.447778624 | 0.03559 |
| ADAMTS18 | -2.453459151 | 0.00422 |
| LINC00634 | -2.458454493 | 0.03953 |
| PSMG3 | -2.470744306 | 0.00702 |
| CDX2 | -2.478733777 | 0.00034 |
| AK098303 | -2.48590786 | 0.03733 |
| NAA11 | -2.493976286 | 0.00005 |
| NEB | -2.526419563 | 0.04379 |
| OR7A5 | -2.532456953 | 0.01883 |
| CALB1 | -2.543602972 | 0.03556 |

| TCGA, BLBC | | |
|---|---|---|
| Gene | logFC | P.Value |
| ALPI | 4.769925 | 0.00191946 |
| PRAMEF8 | 4.70043972 | 0.00392003 |
| SSX2 | 4.67939373 | 0.01316319 |
| OSTN | 4.1337835 | 0.00176187 |
| UGT2B11 | 3.81531009 | 0.02233415 |
| XAGE1D | 3.78646867 | 0.02639049 |
| PABPC1L2B | 3.6701519 | 0.02132082 |
| PTPRT | 3.64983451 | 8.50E-06 |
| OR8A1 | 3.60992113 | 0.00195742 |
| CGA | 3.55726059 | 0.04434046 |
| MAGEB6 | 3.34063247 | 0.00924267 |
| NCRNA00200 | 3.30297421 | 0.01912291 |
| BPESC1 | 3.20943162 | 0.00595498 |
| GPR64 | 3.09924532 | 5.14E-05 |
| PANX3 | 3.08017506 | 0.0059142 |
| HS3ST4 | 3.07527343 | 0.00034294 |
| GABRG2 | 3.05749104 | 0.00239052 |
| ZIC2 | 3.0465538 | 0.00261145 |
| FAM123A | 3.01202226 | 0.00131872 |
| DRD2 | 2.99742808 | 3.08E-06 |
| TP73 | 2.96144635 | 1.50E-07 |
| VIT | 2.94667087 | 8.10E-06 |
| NFASC | 2.92584207 | 7.86E-06 |
| LOC348021 | 2.92492902 | 0.05180449 |
| CYP4X1 | 2.9207511 | 0.0001008 |
| CRTAC1 | 2.87404731 | 0.0002275 |
| ADCY5 | 2.85809697 | 3.28E-05 |
| ZBTB8B | 2.85808806 | 8.16E-05 |
| SSX4 | 2.84121691 | 0.02141746 |
| LEFTY2 | 2.82537633 | 0.000636 |
| KIAA0319 | 2.80204189 | 7.37E-06 |
| FAM135B | 2.74453756 | 0.00033217 |
| NGFR | 2.72813728 | 1.92E-06 |
| BMP7 | 2.71770217 | 0.00054048 |
| ADAMTS18 | 2.68419729 | 2.44E-05 |
| GP2 | 2.68028343 | 0.01533261 |
| FSTL4 | 2.67952329 | 0.00012373 |
| HOXB13 | 2.65050022 | 0.01111597 |
| NTRK2 | 2.6485934 | 1.83E-06 |
| SPSB4 | 2.59502837 | 0.00036131 |
| RET | 2.59075573 | 8.03E-06 |
| DEFB132 | 2.57646585 | 0.02185179 |
| C1QL1 | 2.56560947 | 0.00015401 |
| MYBPC1 | 2.55612213 | 0.01002825 |
| ABCA13 | 2.5477581 | 0.00079225 |
| IGFBPL1 | 2.53874659 | 0.00014519 |
| PEG3 | 2.52592355 | 0.00710329 |
| MYO16 | 2.51022257 | 1.64E-05 |
| KCNB1 | 2.5009036 | 0.00019427 |
| UNC5D | 2.49116825 | 0.0063889 |
| MUC16 | 2.49083659 | 0.00478554 |
| DSCR4 | 2.49020853 | 0.05402415 |
| C10orf90 | 2.48512597 | 0.00055527 |
| TCEAL2 | 2.4755231 | 0.00249308 |
| C2orf40 | 2.47185514 | 0.00268219 |
| HOTAIR | 2.47133479 | 0.00025374 |
| EGF | 2.46034714 | 0.00017737 |
| MGAT5B | 2.46019527 | 0.00015097 |
| SCARA5 | 2.45647702 | 0.00017773 |
| SLC35F3 | 2.45607303 | 0.00071146 |
| C3orf15 | 2.45428124 | 4.07E-05 |
| LGR6 | 2.43062482 | 0.00259813 |
| ACSL6 | 2.42609176 | 0.00054393 |
| AQP5 | 2.4126752 | 0.01414582 |
| CSMD3 | 2.40356486 | 0.00166836 |
| SLC17A8 | 2.39853304 | 0.01252614 |
| ADAM22 | 2.39450391 | 3.24E-05 |
| FABP7 | 2.38690602 | 0.0067096 |
| MSTN | 2.36776749 | 9.52E-07 |
| FCAMR | 2.35728859 | 0.00207052 |
| COL11A2 | 2.3546374 | 0.00059659 |
| ZNF541 | 2.35341686 | 1.39E-06 |
| APLP1 | 2.35165613 | 1.01E-05 |
| PDZD4 | 2.35058517 | 0.00011228 |
| TFF1 | 2.34396247 | 0.04663419 |
| LONRF2 | 2.341803 | 0.00047517 |
| KIRREL2 | 2.34124529 | 0.00076958 |
| 11-Mar | 2.34123976 | 0.01117983 |
| RNF157 | 2.3384221 | 1.46E-07 |
| ALB | 2.33614795 | 0.00103669 |
| NRG3 | 2.32672907 | 0.00124581 |
| DCX | 2.32240806 | 0.01011941 |
| ANGPTL7 | 2.31884339 | 0.01048173 |
| NCAM1 | 2.31289858 | 8.54E-05 |
| CDH4 | 2.31233991 | 0.00213577 |
| SNAP25 | 2.31108231 | 6.53E-06 |
| CEACAM22P | 2.30800314 | 6.99E-05 |
| DACH2 | 2.2951729 | 0.00064429 |
| FIGF | 2.29337546 | 3.07E-05 |
| REEP1 | 2.293073 | 3.68E-05 |
| SORCS3 | 2.28951488 | 0.02494373 |
| TAGLN3 | 2.27539333 | 0.00140709 |

| TCGA, TNBC | | |
|---|---|---|
| Gene | logFC | P.Value |
| ADAM7 | 5.714245518 | 0.03321954 |
| LOC642929 | 4.857980995 | 0.00145424 |
| GAGE2D | 4.810456685 | 0.0383465 |
| ELSPBP1 | 4.169925001 | 0.02105551 |
| SSX2 | 4.005907392 | 0.03025068 |
| GAGE8 | 3.973611276 | 0.03966881 |
| SPACA1 | 3.807354922 | 0.04874737 |
| OR8A1 | 3.709460499 | 0.00316211 |
| CGA | 3.437170756 | 0.00198869 |
| LOC348021 | 3.32492902 | 0.01818799 |
| C7 | 3.170246407 | 5.68E-05 |
| PABPC1L2B | 2.881018425 | 0.01344203 |
| PIGR | 2.853126792 | 0.00173869 |
| GABRG2 | 2.83078452 | 0.03458823 |
| CRTAC1 | 2.813519774 | 1.59E-05 |
| TFF1 | 2.792195812 | 0.01268613 |
| SLC38A8 | 2.757270334 | 0.00919325 |
| SLC44A4 | 2.735312897 | 0.00056991 |
| ANGPTL7 | 2.729189436 | 0.00385304 |
| GAGE2A | 2.725929618 | 0.04538448 |
| OR7E5P | 2.672449623 | 0.03534633 |
| FCAMR | 2.669343994 | 7.34E-05 |
| NCRNA00052 | 2.661039672 | 0.01195538 |
| LY6G6D | 2.624832194 | 0.03215633 |
| NGFR | 2.554816762 | 5.74E-07 |
| PTPRT | 2.552657607 | 0.00027074 |
| ABCA8 | 2.541942223 | 2.03E-05 |
| VIT | 2.525664901 | 0.00022548 |
| DRD2 | 2.514807943 | 2.19E-05 |
| ABCA13 | 2.351888464 | 0.00097296 |
| PEG3 | 2.350533558 | 0.00562794 |
| C1orf146 | 2.321928095 | 0.03256025 |
| CADM3 | 2.315570702 | 0.00011626 |
| SLC7A3 | 2.308138396 | 0.00092039 |
| DSCAML1 | 2.304323823 | 1.46E-05 |
| ALB | 2.282947622 | 0.00016014 |
| KIRREL2 | 2.27486237 | 0.00043675 |
| C20orf114 | 2.262569516 | 0.046725 |
| TP73 | 2.258174427 | 1.87E-05 |
| SERPINA11 | 2.257518278 | 0.00202861 |
| COL11A2 | 2.235717668 | 0.00121666 |
| PLA2G2A | 2.234948466 | 0.00394852 |
| HPCAL4 | 2.226773342 | 2.85E-07 |
| FBN3 | 2.226347371 | 0.00915598 |
| TCEAL2 | 2.209844999 | 0.00321494 |
| BCAN | 2.206988398 | 0.00307728 |
| C3orf15 | 2.200495244 | 3.10E-05 |
| ADH1B | 2.195065528 | 0.02945002 |
| KCNH3 | 2.194414286 | 2.65E-06 |
| LEFTY2 | 2.188631546 | 0.00366389 |
| TMEM59L | 2.171111076 | 3.24E-06 |
| FAM135B | 2.147689477 | 0.0005946 |
| CRLF1 | 2.128669932 | 0.00274967 |
| CMTM5 | 2.122300929 | 0.02677471 |
| KCNC2 | 2.112916521 | 0.03583952 |
| KCNB1 | 2.111357095 | 0.00070605 |
| ADCY5 | 2.109804586 | 0.00027355 |
| MUC4 | 2.109687118 | 0.00087332 |
| ALPPL2 | 2.107448434 | 0.01653342 |
| CD300LG | 2.105907769 | 0.00054585 |
| DARC | 2.104579101 | 8.98E-05 |
| PSD2 | 2.094292737 | 9.25E-06 |
| VSIG2 | 2.074173441 | 0.0041471 |
| SLC13A2 | 2.073157637 | 0.00826831 |
| FIGF | 2.069070848 | 2.42E-05 |
| HS3ST4 | 2.063154337 | 0.01461291 |
| GRPR | 2.054248392 | 0.00014734 |
| PGM5 | 2.049250323 | 3.01E-06 |
| GRIK3 | 2.041626159 | 0.00064521 |
| FABP4 | 2.037073051 | 0.01629332 |
| HLF | 2.034362803 | 1.26E-06 |
| FAM163B | 2.028231216 | 0.00587802 |
| HTR3E | 2.022609093 | 0.00103446 |
| CA10 | 2.020131286 | 0.04210319 |
| PHF21B | 2.016670795 | 0.00336995 |
| ABCA10 | 2.014581087 | 6.35E-06 |
| SLC17A8 | 2.012416243 | 0.00990096 |
| AGR3 | 2.011578159 | 0.02519675 |
| OLFM1 | 2.0070064 | 7.51E-05 |
| OGN | 2.003993588 | 0.00232626 |
| CYP4X1 | 1.986477622 | 0.00209963 |
| MPPED2 | 1.98472198 | 2.92E-05 |
| CPLX3 | 1.975764542 | 0.00400472 |
| TPO | 1.969731233 | 7.04E-06 |
| NTRK2 | 1.969211345 | 0.00043381 |
| TCP10 | 1.968963532 | 0.04247502 |
| RGS9 | 1.96840678 | 0.00067495 |
| CYP4Z1 | 1.954322961 | 0.04180635 |
| KRT78 | -1.999247481 | 0.00325512 |
| OR51B4 | -2.000174484 | 0.01801397 |
| C10orf99 | -2.004527097 | 0.00803887 |

| | | |
|---|---|---|
| CDX2 | -2.401169832 | 0.006452237 |
| SPRR3 | -2.441228869 | 0.024739367 |
| ANKRD53 | -2.453846199 | 0.001878242 |
| GABRQ | -2.501146314 | 0.044903436 |
| KRTAP2 | -2.548637667 | 0.033642949 |
| DTHD1 | -2.565486595 | 0.047570692 |
| CEACAM3 | -2.723405288 | 0.025017413 |
| PKD1L2 | -2.763740872 | 0.007404966 |
| ST6GALNAC1 | -2.831387322 | 0.031461658 |
| PCDHA5 | -2.848343512 | 0.036222013 |
| C5orf58 | -2.855074592 | 0.041675064 |
| NEB | -2.866211337 | 0.015925502 |
| AK055145 | -2.934714236 | 0.008211822 |
| UCN3 | -2.979796716 | 0.049892677 |
| TAS2R20 | -2.997311776 | 0.054397372 |
| LDLRAD1 | -3.080739492 | 0.049392514 |
| ACER1 | -3.129316164 | 0.028042973 |
| VSIG1 | -3.213610381 | 0.011822165 |
| TERT | -3.234428481 | 0.001813266 |
| LOC101929371 | -3.327295945 | 0.028241422 |
| LINC00704 | -3.380455548 | 0.000121024 |
| KLRC2 | -3.418587544 | 0.035159152 |
| LOC100128885 | -3.545123825 | 0.046663413 |
| SLC5A9 | -3.570385843 | 0.010350715 |
| TPTE2 | -3.59066614 | 0.005005009 |
| KCND3 | -3.767100661 | 0.006359175 |
| LIPK | -3.786606068 | 0.018155876 |
| SYCP1 | -4.013518443 | 0.000737165 |
| PTENP1 | -4.026026206 | 0.007491011 |
| HTR1B | -4.099249722 | 0.000673801 |
| BC042048 | -4.252234712 | 0.021526668 |
| FSIP2 | -4.330980777 | 0.006775693 |
| BARHL2 | -4.533341133 | 0.036224616 |
| TRIM61 | -4.824186152 | 0.045914257 |
| PLA2G4D | -4.851159208 | 0.00494489 |
| PSD2 | -5.03177704 | 0.00264492 |
| MIR1287 | -6.807836626 | 0.001615764 |
| MRVI1 | -7.471754307 | 0.001013811 |
| LOC149134 | -9.346958153 | 0.009217588 |
| CTNNA3 | -9.503987925 | 0.001025283 |

| | | |
|---|---|---|
| NKPD1 | -2.546634759 | 0.02303 |
| HPD | -2.549056925 | 0.03248 |
| AK054623 | -2.549610161 | 0.03992 |
| KIAA1045 | -2.564911299 | 0.03499 |
| GQ868703 | -2.58909239 | 0.02489 |
| FGF3 | -2.59115296 | 0.03127 |
| RAB3C | -2.627685687 | 0.01705 |
| GLIPR1L2 | -2.769083613 | 0.03207 |
| NANOGP1 | -2.769676445 | 0.00240 |
| CTB | -2.77607856 | 0.03741 |
| ZNF205 | -2.779595924 | 0.00515 |
| AFF2 | -2.801298499 | 0.04650 |
| SPDYE3 | -2.845122886 | 0.02677 |
| DQ570052 | -2.861320139 | 0.02268 |
| AJ004954 | -2.928448457 | 0.00170 |
| HMX2 | -2.954219402 | 0.01644 |
| BC022047 | -2.983168874 | 0.01477 |
| CBLN1 | -2.996123794 | 0.03118 |
| AK055145 | -3.11419025 | 0.01352 |
| RP11 | -3.145722571 | 0.00621 |
| ZNF541 | -3.150156428 | 0.00349 |
| GAS6 | -3.20021277 | 0.04143 |
| BC112312 | -3.205005731 | 0.01768 |
| NLRP7 | -3.221972591 | 0.03801 |
| OPRD1 | -3.276302988 | 0.02109 |
| MLLT10P1 | -3.32902785 | 0.00832 |
| AX746967 | -3.411509805 | 0.01393 |
| USP41 | -3.536409211 | 0.02375 |
| RAB6C | -3.549294489 | 0.00866 |
| TPTE2 | -3.59066614 | 0.04414 |
| BC015433 | -3.600395762 | 0.03024 |
| KCP | -3.630038375 | 0.02330 |
| LOC101929371 | -3.779123688 | 0.01185 |
| GBX2 | -3.868045023 | 0.00145 |
| TEKT3 | -4.279954838 | 0.01309 |
| INS | -4.327887578 | 0.00222 |
| A2M | -4.328395239 | 0.01081 |
| CEACAM3 | -4.329816963 | 0.04844 |
| KLRC2 | -4.543365862 | 0.02354 |
| UGT2A3 | -4.909014752 | 0.00005 |
| WFDC5 | -4.950341087 | 0.00959 |
| SLC10A4 | -5.59297842 | 0.00606 |
| MEGF11 | -5.652401261 | 0.02145 |
| FGF19 | -5.79777243 | 0.01919 |
| C6orf165 | -5.930897098 | 0.00263 |
| TCRBV13S5 | -6.383193393 | 0.00524 |
| PRSS41 | -6.931128841 | 0.00011 |
| CCDC17 | -7.019501525 | 0.02745 |
| NPFFR1 | -7.812356257 | 0.02420 |

| | | |
|---|---|---|
| LAMA3 | 2.26834126 | 8.19E-06 |
| DKK1 | 2.26454674 | 0.00370069 |
| CNTN2 | 2.26353702 | 0.0002526 |
| TMEM59L | 2.26255183 | 1.51E-05 |
| KCNH3 | 2.26151492 | 8.65E-06 |
| TRIM9 | 2.25569206 | 1.30E-05 |
| UGT2B15 | 2.25447906 | 0.00578208 |
| MPPED2 | 2.25353199 | 8.87E-05 |
| TRPM6 | 2.25316513 | 0.00056613 |
| EPHA10 | 2.25203238 | 2.55E-05 |
| NALCN | 2.22742649 | 0.00067743 |
| C1orf94 | 2.22193896 | 0.03469977 |
| CRLF1 | 2.21909243 | 0.00822098 |
| PPP1R1A | 2.21105315 | 0.0057855 |
| AGR3 | 2.2087265 | 0.02547382 |
| KGFLP1 | 2.20705129 | 4.03E-05 |
| BPIL3 | 2.19615871 | 0.03740995 |
| RELN | 2.19497413 | 0.00095133 |
| MYRIP | 2.19429425 | 0.00035178 |
| CNTFR | 2.19336756 | 0.00715889 |
| BCAN | 2.18998729 | 0.01191845 |
| GKN2 | 2.18738401 | 0.001227 |
| PAK7 | 2.18548205 | 0.00010519 |
| ATP4B | 2.18438628 | 0.00049823 |
| LRRN2 | 2.17953256 | 9.22E-05 |
| AGXT | 2.1792395 | 0.04670918 |
| CNTN5 | 2.16892078 | 0.0048458 |
| LYPD6 | 2.16662767 | 2.91E-05 |
| ZBTB16 | 2.16609198 | 0.00034971 |
| SYT13 | 2.16583311 | 0.0261753 |
| CD300LG | 2.16443805 | 0.00461307 |
| FLJ12825 | 2.16374654 | 2.95E-05 |
| ALX4 | 2.14540075 | 0.00790716 |
| JAKMIP1 | 2.14070524 | 0.00026679 |
| C4BPA | 2.13922351 | 0.00670343 |
| LHX2 | 2.13830816 | 0.00093837 |
| PHF21B | 2.13249013 | 0.00550671 |
| KSR2 | 2.11876042 | 0.00040847 |
| ATRNL1 | 2.11314974 | 0.0077298 |
| RGS9 | 2.09671095 | 0.00244697 |
| ABCA10 | 2.08971301 | 4.75E-06 |
| ODZ2 | 2.08772057 | 0.00040929 |
| USP44 | 2.08068645 | 0.00021401 |
| FRMPD4 | 2.07884547 | 0.00282057 |
| MATN2 | 2.07706237 | 3.09E-05 |
| CMTM5 | 2.06633074 | 0.03014593 |
| TMPRSS5 | 2.0619219 | 0.00079965 |
| TFF3 | 2.06031957 | 0.02679928 |
| SCUBE2 | 2.05956914 | 0.00016425 |
| LOC10019093 | 2.05765454 | 3.78E-05 |
| MEGF10 | 2.05489027 | 0.00099338 |
| CHRDL2 | 2.05247932 | 0.0096111 |
| GPR143 | 2.04964277 | 0.00015707 |
| CCDC129 | 2.0458384 | 0.00883065 |
| XIST | 2.04569904 | 5.01E-06 |
| ALDH8A1 | 2.03645167 | 1.49E-06 |
| SLC22A3 | 2.02000602 | 0.00041539 |
| CACNB4 | 2.00331001 | 0.00017743 |
| PAK3 | 2.00313111 | 0.0002573 |
| PSD2 | 1.99954317 | 0.00065213 |
| HTR3E | 1.99835795 | 0.0012338 |
| SOX8 | 1.99514575 | 0.01273639 |
| S100P | -1.9614501 | 0.02631588 |
| KRT4 | -2.0020207 | 0.04588919 |
| LCN2 | -2.1019715 | 0.00842713 |
| CWH43 | -2.1232815 | 0.01570089 |
| ADH7 | -2.1415763 | 0.05358521 |
| ZCCHC16 | -2.1756875 | 0.01463716 |
| KRT79 | -2.2267249 | 0.02585675 |
| PSAPL1 | -2.2332604 | 0.03963001 |
| TDRD1 | -2.2892698 | 0.02376192 |
| SPRR1A | -2.2925772 | 0.01926323 |
| CA9 | -2.4991546 | 0.00119703 |
| LCE2A | -2.6297223 | 0.01631921 |
| LY6D | -2.631763 | 0.00681274 |
| OC90 | -2.6996269 | 0.04516405 |
| VSTM2B | -2.9321935 | 0.04889516 |
| S100A7 | -3.0333387 | 0.04802538 |
| DPP10 | -3.1546539 | 0.00503713 |
| PRAMEF1 | -3.169925 | 0.00428433 |
| MT4 | -3.3219281 | 0.04564901 |
| C10orf129 | -3.6892849 | 2.89E-06 |
| RETNLB | -3.7004397 | 0.01522475 |
| SEMG2 | -4.2637385 | 0.02098346 |
| DNAJC5G | -5.1978951 | 3.22E-05 |
| GJA8 | -5.4594316 | 4.54E-05 |
| FTHL17 | -5.9985009 | 0.00500081 |
| NXF2 | -8.4426696 | 0.00020272 |

| | | |
|---|---|---|
| EPS8L3 | -2.009043403 | 0.02682371 |
| FAM25A | -2.031415226 | 0.04227349 |
| FOXG1 | -2.057088689 | 0.04731655 |
| TNP1 | -2.069800224 | 0.04517713 |
| CSRP3 | -2.163951032 | 0.04043304 |
| S100A8 | -2.213164841 | 0.00232077 |
| PLA2G2F | -2.227270831 | 0.0184851 |
| MMP1 | -2.247894759 | 0.00145372 |
| CNBD1 | -2.299309881 | 0.00964119 |
| LY6D | -2.32050497 | 0.00640054 |
| SPANXA2 | -2.395383247 | 0.04964038 |
| C10orf129 | -2.395843219 | 0.01357891 |
| DGAT2L6 | -2.464385619 | 0.03097603 |
| TMEM8C | -2.471679166 | 0.03811362 |
| KRT79 | -2.508123781 | 0.00695036 |
| DPP10 | -2.648165149 | 0.01644482 |
| RPTN | -2.876242708 | 0.02371727 |
| CA9 | -2.894237811 | 0.00016249 |
| C17orf77 | -2.895891196 | 0.00960288 |
| HTR3B | -3.272896374 | 0.05281384 |
| S100A7A | -3.306489796 | 0.01346404 |
| F9 | -3.564641508 | 0.0383062 |
| MT4 | -3.704695468 | 0.00434049 |
| S100A7 | -3.73998807 | 0.0037761 |
| PTPN20A | -4.279735821 | 0.03365632 |
| SPINT3 | -4.321928095 | 0.0243849 |
| PMCHL2 | -4.33974005 | 0.04461753 |
| CFHR5 | -4.690492758 | 0.0022544 |
| VSTM2B | -4.857980995 | 0.02629207 |
| PNLIP | -5.166577675 | 0.00124963 |
| GJA8 | -5.459431619 | 0.0013155 |
| ISX | -5.873288558 | 0.0096147 |
| NXF2 | -7.633624223 | 0.00017764 |

Supplemental Fig. 3

a.

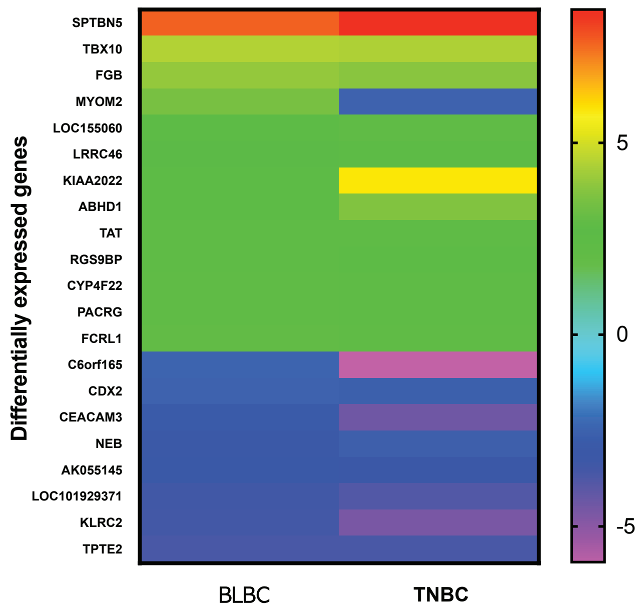

b.

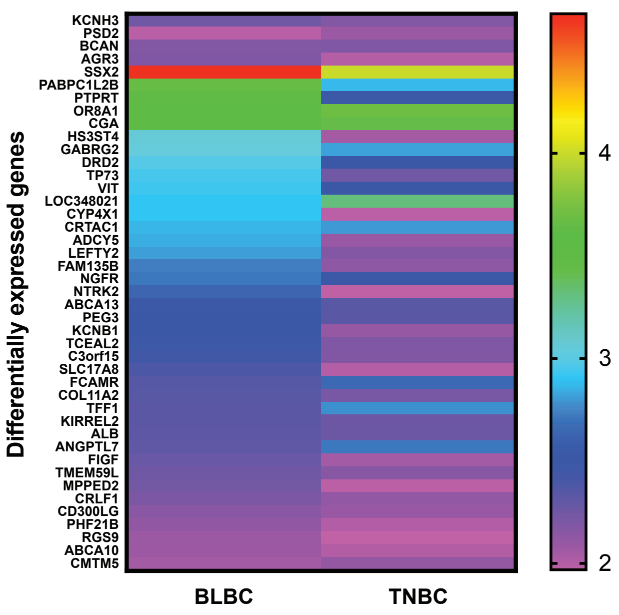

c.

BLBC and TNBC patient overlap

SCANB, large data set

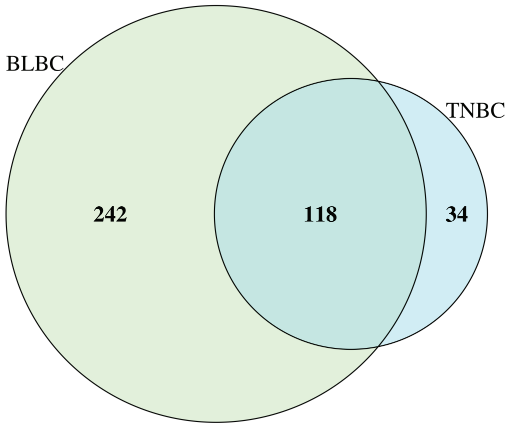

d.

BLBC and TNBC patient overlap

TCGA

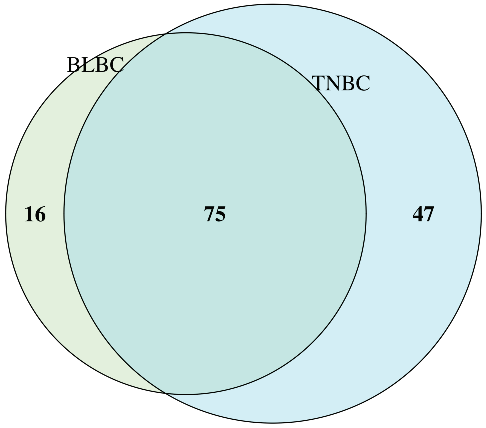

e.

| SCAN-B, BLBC | | | |
| --- | --- | --- | --- |
| | Motif | E-value | Unerased E-value |
| 1 | AAATAYAA | 4.00E-17 | 4.00E-17 |
| 2 | GTAGAGAY | 8.40E-17 | 8.40E-17 |
| 3 | GGATTAYA | 1.00E-15 | 1.00E-15 |
| 4 | GAGGCTGA | 1.30E-14 | 1.30E-14 |
| 5 | AAAAAAAA | 1.60E-13 | 1.60E-13 |
| 6 | AGTAGCTG | 2.20E-13 | 2.20E-13 |
| 7 | TCACYGCA | 3.00E-13 | 3.00E-13 |
| 8 | ACTGYACT | 7.20E-13 | 7.20E-13 |
| 9 | CAGCCTGG | 2.30E-12 | 2.30E-12 |
| 10 | CACCRCRC | 1.90E-10 | 7.60E-11 |
| 11 | GCTCACRC | 2.60E-10 | 2.60E-10 |
| 12 | CCKCCRCC | 4.60E-10 | 4.60E-10 |
| 13 | GTCTCGAW | 1.20E-09 | 4.50E-10 |
| 14 | GASACAGA | 8.10E-10 | 2.70E-09 |
| 15 | CAAAGTGC | 1.50E-09 | 1.50E-09 |
| 16 | AAATTAGC | 1.60E-08 | 7.70E-10 |
| 17 | GCAGGAGR | 2.40E-07 | 2.40E-07 |
| 18 | GGTCAGGM | 4.80E-07 | 4.80E-07 |
| 19 | TATTTWTW | 1.50E-06 | 1.80E-11 |
| 20 | CYGGGYTC | 2.30E-06 | 1.90E-06 |
| 21 | CTGTARTC | 3.90E-06 | 1.30E-12 |
| 22 | GBGACAGA | 4.70E-06 | 6.90E-07 |
| 23 | CACCCGCS | 8.00E-06 | 6.80E-04 |
| 24 | GGTGAAAC | 2.50E-05 | 2.50E-05 |
| 25 | GAGGCMGA | 7.40E-05 | 7.40E-05 |
| 26 | CACYGTGC | 8.70E-05 | 8.70E-05 |
| 27 | TTTTAAAA | 8.70E-05 | 2.40E-04 |
| 28 | GAGHGAGA | 1.60E-04 | 9.50E-06 |
| 29 | GTGTGTGY | 1.90E-04 | 5.50E-04 |
| 30 | GAGAKCGC | 2.70E-04 | 2.70E-04 |
| 31 | ARGTGATC | 7.50E-04 | 3.50E-04 |
| 32 | ACCATGCC | 1.30E-03 | 1.30E-03 |
| 33 | CACSTGCC | 1.80E-03 | 4.10E-04 |
| 34 | AKWAAATA | 3.70E-03 | 2.50E-04 |
| 35 | CCCRCCYC | 3.80E-03 | 3.90E-05 |
| 36 | GAGAYGGA | 2.30E-03 | 8.30E-05 |
| 37 | CGCSGCSC | 8.80E-03 | 3.80E-04 |
| 38 | AAAAAYAA | 2.00E-02 | 1.90E-02 |
| 39 | GAAATAAK | 2.40E-02 | 1.20E-02 |

| SCAN-B, TNBC | | | |
| --- | --- | --- | --- |
| | Motif | E-value | Unerased E-value |
| 1 | TAMAAATA | 2.20E-15 | 2.20E-15 |
| 2 | CACKGCAC | 3.20E-13 | 3.20E-13 |
| 3 | CRGCCTCC | 6.50E-12 | 6.50E-12 |
| 4 | AAAAAWAA | 1.80E-11 | 2.90E-11 |
| 5 | CTGTAATC | 2.90E-11 | 2.90E-11 |
| 6 | AGGCTGGA | 1.00E-09 | 4.20E-09 |
| 7 | CCCAGCTA | 1.90E-09 | 2.30E-10 |
| 8 | GAGRCGGA | 3.50E-09 | 3.50E-09 |
| 9 | GCCYGGCC | 4.70E-09 | 1.50E-09 |
| 10 | RAGACAGA | 4.50E-08 | 4.50E-08 |
| 11 | AYATATRT | 1.20E-07 | 1.20E-07 |
| 12 | AAAAAWTA | 2.20E-07 | 5.90E-08 |
| 13 | AAATAAAH | 2.70E-07 | 1.10E-07 |
| 14 | CAGTGARC | 3.80E-07 | 3.80E-07 |
| 15 | CCACCDCC | 1.10E-06 | 3.80E-07 |
| 21 | CACCRYGC | 1.80E-06 | 1.10E-07 |
| 16 | CTCCTGAC | 6.00E-06 | 9.90E-07 |
| 17 | CGCVCGCC | 7.70E-06 | 7.70E-06 |
| 18 | TGGWGAAA | 8.60E-06 | 3.80E-07 |
| 19 | AGGCAGGA | 8.90E-06 | 8.90E-06 |
| 20 | SGAGACCA | 9.90E-06 | 2.00E-05 |
| 22 | ATCGCTTG | 1.80E-05 | 1.80E-05 |
| 23 | CWCACAC | 2.40E-05 | 2.40E-05 |
| 24 | AAGTGCTG | 3.50E-05 | 3.50E-05 |
| 25 | AMAGAAAA | 5.40E-05 | 1.20E-04 |
| 26 | CABAGTGA | 5.80E-05 | 5.80E-05 |
| 27 | GTAGAGAY | 6.50E-05 | 2.80E-05 |
| 28 | ATTWTAWA | 1.30E-04 | 4.60E-06 |
| 30 | GCGTGAGC | 1.40E-04 | 7.90E-04 |
| 29 | CACCRCAC | 3.80E-04 | 1.40E-05 |
| 31 | GCGGCGCB | 1.20E-03 | 5.10E-04 |
| 32 | TCACTTKA | 1.30E-03 | 1.30E-03 |
| 33 | CTGCCKCC | 1.60E-03 | 1.40E-04 |
| 34 | ACTACAGG | 3.80E-03 | 5.90E-04 |
| 35 | CCAGCTVC | 6.10E-03 | 1.00E-04 |
| 36 | CTGGYTAA | 6.20E-03 | 1.50E-04 |
| 37 | GAAAAAKA | 7.30E-03 | 3.40E-04 |
| 38 | CTCCCAYC | 1.00E-02 | 6.50E-04 |
| 39 | CTGTCRCC | 1.70E-02 | 1.00E-04 |
| 40 | CCGCDGCC | 1.70E-02 | 3.30E-02 |

Supplemental Fig. 4

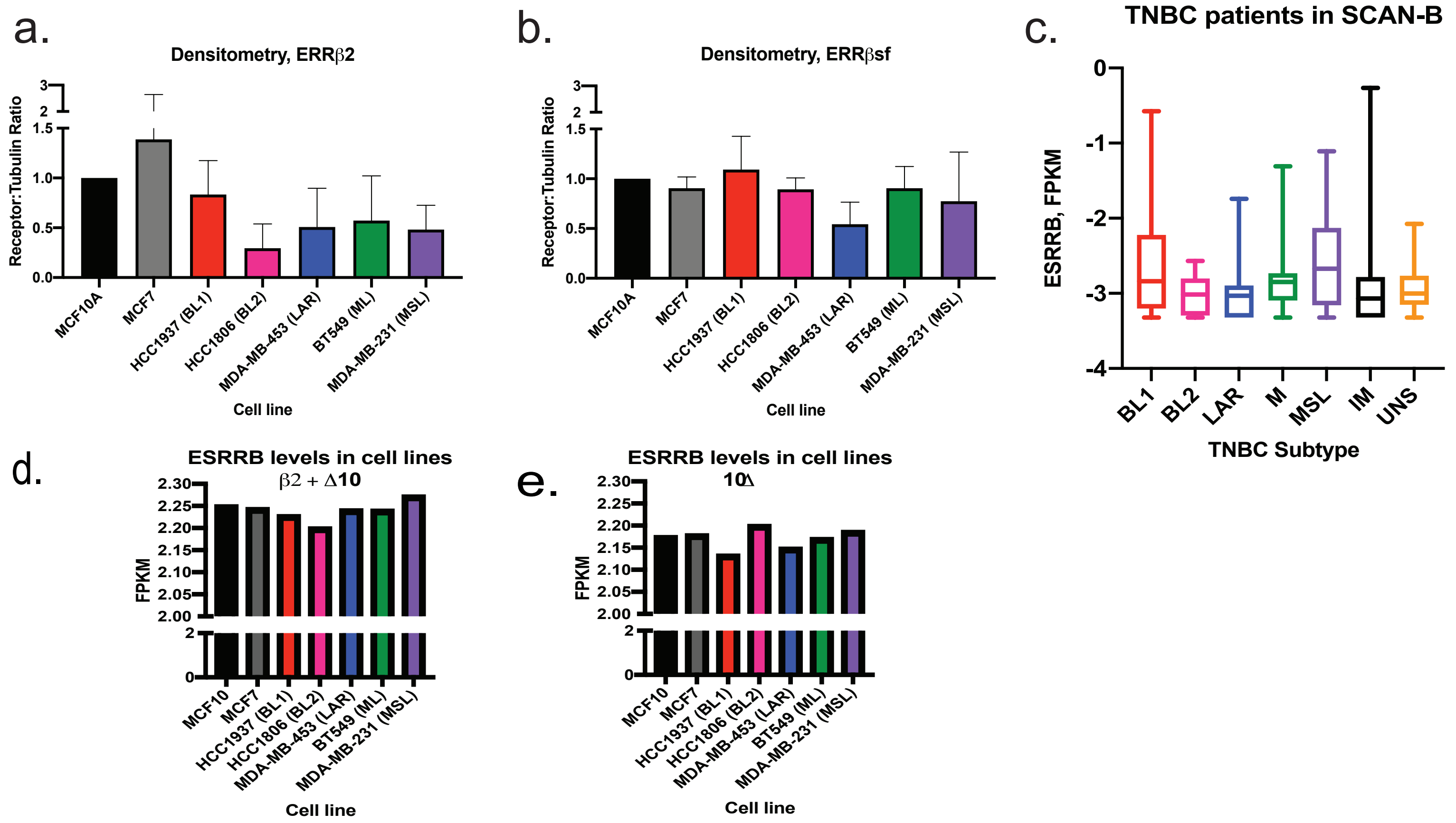

Supplemental Table 2

2.1

| | ERRβ2 | ERRβsf | ERRβ2:ERRβsf |
|---|---|---|---|
| | Mean(sd), median[IQR] | | |
| **ER+ (n=50)** | 0.86(0.10), 0.87[0.82, 0.94] | 0.66(0.16), 0.68[0.58, 0.77] | 1.39(0.52), 1.27[1.11, 0.49] |
| **HER2+ (n=50)** | 0.90(0.13), 0.95[0.87, 0.98] | 0.70(0.17), 0.71[0.59, 0.82] | 1.36(0.39), 1.28[1.13, 1.45] |
| **TNBC (n=50)** | 0.85(0.23), 0.94[0.82, 0.98] | 0.71(0.23), 0.77[0.65, 0.88] | 1.28(0.53), 1.17[1.07, 1.30] |
| **Kruskal-Wallis *p*-value** | 0.016* | 0.086 | 0.095 |

2.2

| | ERRβ2 | | ERRβsf | |
|---|---|---|---|---|
| | Nuclear | Cytoplasmic | Nuclear | Cytoplasmic |
| | Mean(sd), median[IQR] | | Mean(sd), median[IQR] | |
| **ER+ (n=50)** | 0.65 (0.36), 0.67 [0.37, 1.00] | 0.55 (0.40), 0.47 [0.16, 0.96] | 0.17 (0.15), 0.13 [0.03, 0.29] | 0.11 (0.09), 0.09 [0.04, 0.14] |
| **HER2+ (n=50)** | 0.98 (0.05), 1.00 [0.99, 1.00] | 0.03 (0.06), 0.01 [0.00, 0.03] | 0.35 (0.25), 0.37 [0.13, 0.46] | 0.20 (0.18), 0.16 [0.07, 0.29] |
| **TNBC (n=50)** | 0.96 (0.17), 1.00 [1.00, 1.00] | 0.07 (0.19), 0.01 [0.00, 0.02] | 0.33 (0.25), 0.32 [0.11, 0.43] | 0.22 (0.20), 0.15 [0.07, 0.31] |

Supplemental Table 3

### ERRβ2

#### Receptor Status and Lymph Node Status

| | Source | DF | Anova SS | Mean Square | F Value | Pr > F |
|---|---|---|---|---|---|---|
| w/ interaction | Receptor Status | 2 | 15.34 | 7.67 | 3.03 | 0.052 |
| | Lymph Node Status | 1 | 1.26 | 1.26 | 0.5 | 0.482 |
| | Receptor Status*Lymph Node Status | 2 | 0.8 | 0.4 | 0.16 | 0.856 |
| w/o interaction | **Source** | **DF** | **Anova SS** | **Mean Square** | **F Value** | **Pr > F** |
| | Receptor Status | 2 | 15.34 | 7.67 | 3.03 | 0.052 |
| | Lymph Node Status | 1 | 1.26 | 1.26 | 0.5 | 0.482 |

#### Receptor Status and Race

| | Source | DF | Anova SS | Mean Square | F Value | Pr > F |
|---|---|---|---|---|---|---|
| w/ interaction | Receptor Status | 2 | 14.55 | 7.28 | 2.88 | 0.059 |
| | Race | 4 | 29.45 | 7.36 | 2.92 | 0.024 |
| | Receptor Status*Race | 6 | 19.08 | 3.18 | 1.27 | 0.27 |
| w/o interaction | **Source** | **DF** | **Anova SS** | **Mean Square** | **F Value** | **Pr > F** |
| | Receptor Status | 2 | 14.55 | 7.28 | 2.88 | 0.059 |
| | Race | 4 | 29.45 | 7.36 | 2.92 | 0.024 |

#### Receptor Status and Age (years)

| | Source | DF | Type III SS | Mean Square | F Value | Pr > F |
|---|---|---|---|---|---|---|
| w/ interaction | Receptor Status | 2 | 0.72496893 | 0.36248447 | 0.13 | 0.8753 |
| | Age (years) | 1 | 1.09642755 | 1.09642755 | 0.4 | 0.5264 |
| | Receptor Status*Age | 2 | 0.26823291 | 0.13411645 | 0.05 | 0.9519 |
| w/o interaction | **Source** | **DF** | **Type III SS** | **Mean Square** | **F Value** | **Pr > F** |
| | Receptor Status | 2 | 12.56 | 6.28 | 2.34 | 0.1 |
| | Age (years) | 1 | 1.13 | 1.13 | 0.42 | 0.518 |

### ERRβsf

#### Receptor Status and Lymph Node Status

| | Source | DF | Anova SS | Mean Square | F Value | Pr > F |
|---|---|---|---|---|---|---|
| w/ interaction | Receptor Status | 2 | 5.63 | 2.82 | 2.41 | 0.094 |
| | Lymph Node Status | 1 | 2.72 | 2.72 | 2.32 | 0.13 |
| | Receptor Status*Lymph Node Status | 2 | 0.71 | 0.35 | 0.3 | 0.741 |
| w/o interaction | **Source** | **DF** | **Anova SS** | **Mean Square** | **F Value** | **Pr > F** |
| | Receptor Status | 2 | 5.63 | 2.82 | 2.41 | 0.094 |
| | Lymph Node Status | 1 | 2.72 | 2.72 | 2.32 | 0.13 |

#### Receptor Status and Race

| | Source | DF | Anova SS | Mean Square | F Value | Pr > F |
|---|---|---|---|---|---|---|
| w/ interaction | Receptor Status | 2 | 4.22 | 2.11 | 1.95 | 0.147 |
| | Race | 4 | 6.93 | 1.73 | 1.6 | 0.179 |
| | Receptor Status*Race | 6 | 19.46 | 3.24 | 2.99 | 0.009 |

#### Receptor Status and Age (years)

| | Source | DF | Type III SS | Mean Square | F Value | Pr > F |
|---|---|---|---|---|---|---|
| w/ interaction | Receptor Status | 2 | 0.37197751 | 0.18598875 | 0.15 | 0.859 |
| | Age (years) | 1 | 0.14527096 | 0.14527096 | 0.12 | 0.7308 |
| | Receptor Status*Age | 2 | 0.14740298 | 0.07370149 | 0.06 | 0.9415 |
| w/o interaction | **Source** | **DF** | **Type III SS** | **Mean Square** | **F Value** | **Pr > F** |
| | Receptor Status | 2 | 3.7 | 1.85 | 1.54 | 0.219 |
| | Age (years) | 1 | 0.18 | 0.18 | 0.15 | 0.699 |
